# Deacidification of distal lysosomes by neuronal aging drives synapse loss

**DOI:** 10.1101/2022.10.12.511921

**Authors:** Tatiana Burrinha, César Cunha, Cláudia Guimas Almeida

## Abstract

Previously, we found that age-dependent beta-amyloid accumulation is not enough to cause synaptic decline. Here, we characterized endolysosomes (late-endosomes and lysosomes) in aged neurons and the aged brain, which might drive synaptic decline since lysosomes are a cellular aging target and relevant for synapses. Neuronal aging induces enlarged endolysosome accumulation in the aged neurons and brain, especially distally, related to the increased anterograde movement. Aged lysosomes abound in neurites but are less degradative due to deacidification despite cathepsin D buildup, leading to late-endosome accumulation. Increasing the acidification of aged lysosomes by ML-SA1 treatment increased degradation and reverted synaptic decline, while lysosome alkalinization by chloroquine treatment mimicked age-dependent lysosome dysfunction and synaptic decline. We identify the deacidification of distal lysosomes as a neuronal mechanism of age-dependent synapse loss. Our findings suggest that future therapeutic strategies to address lysosomal defects might be able to delay age-related synaptic decline.

**Highlights:**

- Enlarged endolysosomes accumulate close to synapses in aged neurons and aged brain
- Late-endosomes accumulate with neuronal aging
- Aged lysosomes are less acidic and degradative despite accumulating Cathepsin D
- Increasing acidification of aged lysosomes improves synapses, while deacidification recapitulates age-dependent synapse loss

**Summary:** We identify the downregulation of the lysosome degradative activity via deacidification as a neuronal aging mechanism contributing to aging-dependent synapse loss.

**Graphical abstract:** 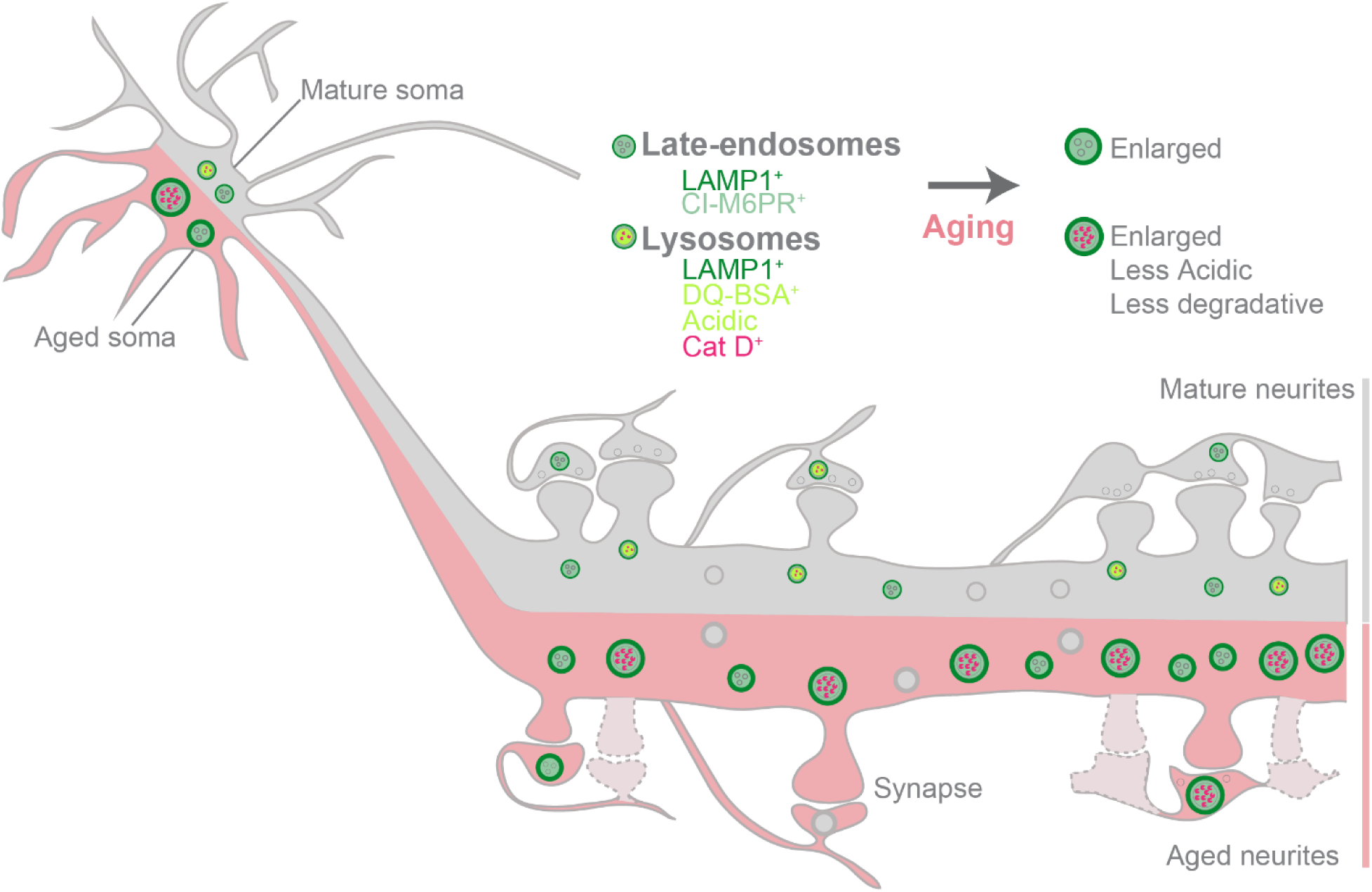

## Introduction

As we age, our functional capabilities decline, and our cells start showing hallmarks of aging (Mattson and Arumugam, 2018), increasing the risk of disease. Indeed, aging is the most significant risk factor for late-onset Alzheimer’s disease (LOAD), the most common neurodegenerative disease of the elderly (Yankner et al., 2008). The deterioration of brain cognitive performance with aging is associated with alterations in the central nervous system, predominantly in the prefrontal cortex and hippocampus (Leal and Yassa, 2015; Yankner et al., 2008), two brain regions with a major role in memory and learning. However, little is known about the cellular and molecular mechanisms underlying cognitive decline during normal brain aging.

At the cellular level, dysfunctional remodeling of postsynaptic compartments, also known as spines, and synapse loss characterize the aging brain and underlie cognitive decline (Dickstein et al., 2013; Morrison and Baxter, 2012; Petralia et al., 2014; Samson and Barnes, 2013). One of the AD earliest mechanisms is synapse loss, occurring before neuronal degeneration, amyloid-beta (Aβ) plaques formation, and tau neurofibrillary tangles (Almeida et al., 2005; Mucke and Selkoe, 2012; Palop and Mucke, 2010; Selkoe and Hardy, 2016). We have shown that *in vitro* neuronal aging potentiates the intraneuronal accumulation of Aβ due to increased endocytosis of amyloid precursor protein (APP). We established causality between Aβ accumulation and synaptic loss in aged neurons. However, the inhibition of Aβ production in aged neurons was insufficient to protect neurons from synapse loss (Burrinha et al., 2021), leading us to postulate that other mechanisms might contribute to age-dependent synapse decline.

Endocytic trafficking increases with aging (Alsaqati et al., 2018; Blanpied et al., 2003; Burrinha et al., 2021). Upon endocytosis, cargo for degradation follows the endo-lysosomal pathway, which comprises endosomes and lysosomes (endolysosomes or ELs). In neurons, endosomes move towards the cell body, maturing into multi-vesicular bodies (MVBs)/ late-endosomes (LEs), which fuse with degradative lysosomes (Ferguson, 2018; Ferguson, 2019; Winckler et al., 2018). The neuronal cell body is considered the primary site of lysosomal degradation (Cai et al., 2010; Cheng et al., 2018; Ferguson, 2019; Gowrishankar et al., 2015; Lee et al., 2011; Tammineni et al., 2017; Yap et al., 2018). In addition, lysosomes degrade autophagic cargo and regulate nutrient sensing, plasma membrane repair, and ion homeostasis (Ballabio and Bonifacino, 2020).

Aging leads to the accumulation of undegradable cellular waste, lipofuscin, such as oxidatively damaged proteins and lipids, in lysosomes, especially in post-mitotic cells like neurons, since it cannot be removed from cells by exocytosis or diluted by cell division (Brunk and Terman, 2002; Gray and Woulfe, 2005; Mattson and Magnus, 2006). Indeed, lysosome dysfunction is an aging mechanism (López-Otín et al., 2013).

Interestingly, lipofuscin, the hallmark of cellular aging, builds up in the aging brain of rodents and humans (Gray and Woulfe, 2005; Sohal and Brunk, 1989; Sohal and Wolfe, 1986) and accumulates in the somatic lysosomes (Burrinha et al., 2021). Also, mRNA and protein expression of Cathepsin D (Cat D), the major lysosomal hydrolase, increased in the aged rat and human brain (Cataldo et al., 1995; Jung et al., 1999; Nakanishi et al., 1994; Nakanishi et al., 1997) suggesting lysosome dysfunction in the physiological aged brain.

Moreover, chemically-induced lysosomal dysfunction causes postsynaptic defects (Goo et al., 2017; Kanju et al., 2007), and genetically induced lysosomal dysfunction drives presynaptic defects in synaptic vesicle recycling (Sambri et al., 2017). Consequently, synapses may be particularly affected by lysosomal dysfunction with aging. Still, it is unknown whether the lysosome becomes dysfunctional with neuronal aging, leading to synapse loss. In this study, we investigated whether ELs’ size, positioning, composition, and function were changed in aged neurons and contributed to synapse loss. We also assessed whether restoring or disrupting lysosome function rescued or induced synapse loss.

We used physiologically aged mouse brains and primary mouse cortical neurons aged for 28 days *in vitro* (DIV). We characterized Els and found that both late-endosomes and lysosomes altered in the soma and distal neurites with aging. In aged neurons, lysosomal degradation was reduced due to deacidification despite cathepsin D buildup, leading to late-endosome accumulation. Finally, we established causality between aging-dependent lysosome dysfunction and synapse decline by restoring lysosome function in aged neurons or inducing lysosome dysfunction in mature neurons. Overall, we implicate lysosomal dysfunction as a causal mechanism of age-dependent synapse loss.

## Results

### Endolysosomes are polarized towards neurites in mature neurons and upregulated with aging

Since lysosome dysfunction is a mechanism of aging (López-Otín et al., 2013), we investigated the impact of neuronal aging on lysosomes. We previously established a cellular aging model of post-mitotic neurons without neurodegeneration (Burrinha et al., 2021), similar to several groups which reported that neurons start aging or cellular senescence after 28 DIV (Aksenova et al., 1999; Goslin and Banker, 1989; Martin et al., 2011; Papa et al., 1995; Trovò et al., 2013). We observed that primary neurons cultured for 21 and 28 DIV did not present gross morphological changes such as axonal bead-like degeneration or dendritic shrinkage, had a residual expression of doublecortin, a neuronal precursor marker, and unaltered expression GFAP, a glia marker (Burrinha et al., 2021). Importantly, 28 DIV neurons showed canonical markers of cellular aging, such as the accumulation of senescence-associated β-galactosidase positive cells (SA-β-Gal) and the formation of the auto-fluorescent granules within lysosomes in cell bodies (Fig. S1 A) (Bigagli et al., 2016; Burrinha et al., 2021; Dong et al., 2014; Gray and Woulfe, 2005; Jurk et al., 2012). This cellular aging of cultured primary neurons likely results from the culture conditions and mimics neuronal changes induced by physiological aging.

We analyzed the distribution of lysosomal-associated protein 1 (LAMP1) labeled compartments that consist of late-endosomes (LEs) and lysosomes (endolysosomes, or ELs)(Cheng et al., 2018; Winckler et al., 2018) in microtubule-associated protein 2 (MAP2)-labeled dendrites of mature and aged neurons. We decided to refer to neurites hereafter instead of dendrites since some LAMP1^+^ ELs may be in the axons that line up with dendrites in these mature cultures. We observed LAMP1^+^ ELs in the cell body/soma and frequently in neurites (Fig. 1 A). We measured the area of LAMP1^+^ ELs in the cell body and the neurites of mature neurons, and we found that the LAMP1^+^ ELs area in neurites was 2-fold larger than in the soma of mature neurons (Fig. 1 B). In aged neurons, the presence of LAMP1^+^ ELs was even more notorious. The endo-lysosomal area increased in aged neurites (48%), indicating that ELs are positioned more distally in aged neurons. Given this relevant alteration in neurites, we next characterized ELs in the soma (somatic ELs) separately from neurites (neuritic ELs).

**Figure 1.**
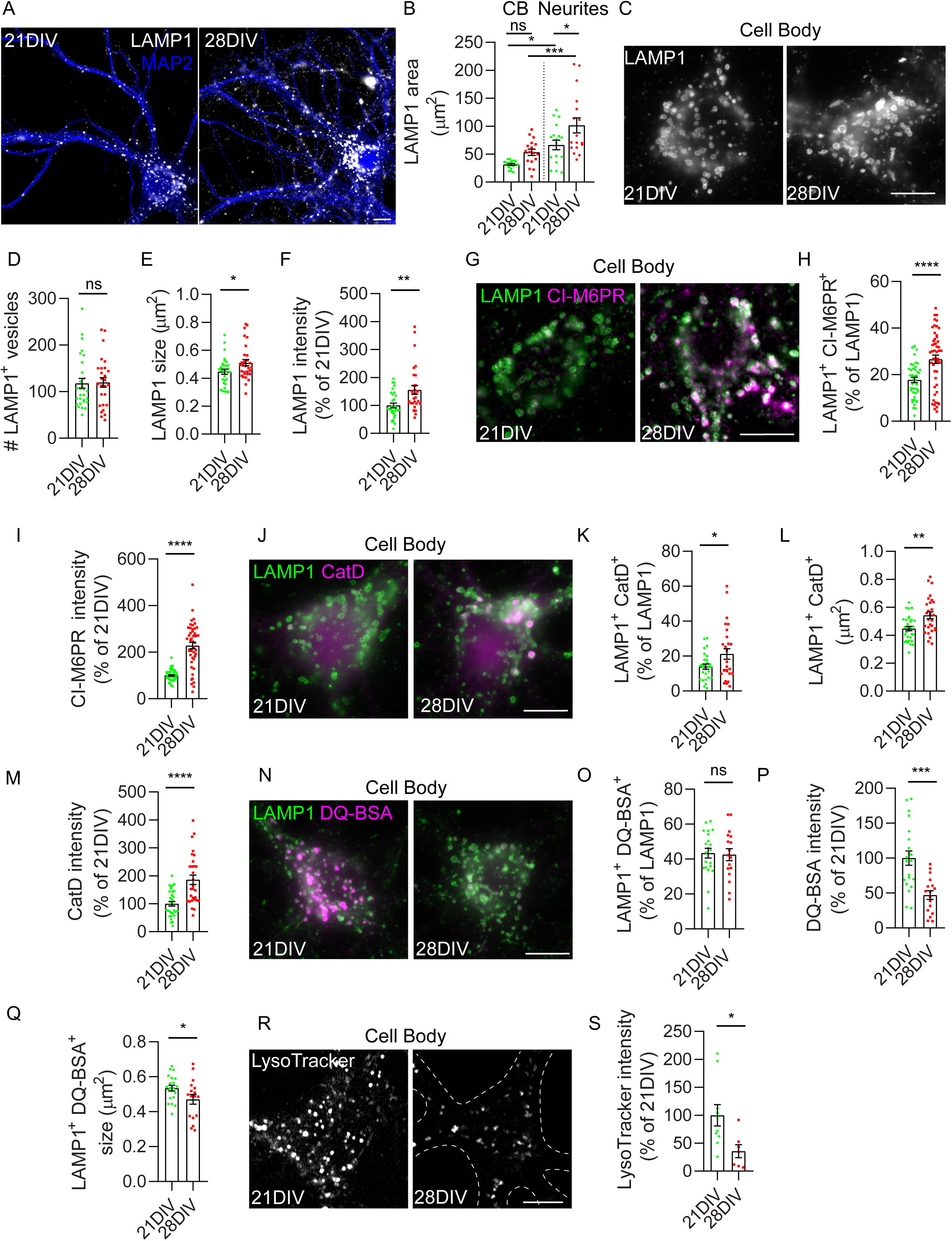
Endolysosomes are upregulated in aged neurons with somatic lysosomes less degradative. (A) Representative image of LAMP1^+^ endolysosomes distribution (white) in mature (21 DIV) and aged (28 DIV) neurons immunolabeled with MAP2 (blue). (B) Quantification LAMP1^+^ area occupied in cell body and neurites of 21 and 28 DIV neurons (n=3, N cell body= 17; N neurites= 17). (C) LAMP1^+^ endolysosomes in the cell body of 21 and 28 DIV neurons. (D) Quantification of LAMP1^+^ endolysosome number in the cell body (n=4, N cell body= 27-29). (E) Quantification of LAMP1^+^ endolysosome size in the cell body (n=4, N cell body= 30-31). (F) Quantification of the mean intensity of LAMP1 *per* endolysosome in the cell body in the percentage of 21 DIV neurons (n=4, N cell body = 29-31). (G) CI-M6PR^+^ (magenta) LAMP1^+^ (green) late-endosomes in the cell body of 21 and 28 DIV neurons, displayed after background subtraction. (H) Quantification of the percentage of LAMP1^+^ endolysosomes colocalizing with CI-M6PR in the cell body (n=3, N cell body= 45-50). (I) Quantification of the mean intensity of CI-M6PR *per* LAMP1^+^ late-endosome in the cell body in the percentage of 21 DIV neurons (n=3, N cell body = 46-56). (J) Cat D^+^ (magenta) LAMP1^+^ (green) lysosomes in the cell body of 21 and 28 DIV neurons. (K) Quantification of the percentage of LAMP1^+^ endolysosomes colocalizing with Cat D in the cell body (n=4, N cell body= 27-28). (L) Quantification of LAMP1^+^ Cat D^+^ lysosomes size in cell body (n=4, N cell body= 28-31). (M) Quantification of the mean intensity of Cat D *per* LAMP1^+^ lysosome in the cell body in the percentage of 21 DIV neurons (n=4, N cell body= 29-32). (N) DQ-BSA^+^ (magenta) LAMP1^+^ (green) lysosomes in the cell body of 21 and 28 DIV neurons. (O) Quantification of the percentage of LAMP1^+^ endolysosomes colocalizing with DQ-BSA in the cell body (n=3, N cell body= 17-21). (P) Quantification of the mean intensity of DQ-BSA *per* LAMP1^+^ lysosome in the cell body in the percentage of 21 DIV neurons (n=3, N cell body= 17-21). (Q) Quantification of LAMP1^+^ DQ-BSA^+^ lysosomes size in cell body (n=3, N cell body= 17-21). (R) LysoTracker puncta in the cell body of 21 and 28 DIV neurons. (S) Quantification of the mean intensity of LysoTracker puncta *per* cell body in the percentage of 21 DIV neurons (n=2, N cell body= 7-10). Data are mean ± SEM. Statistical significance was determined by unpaired t-test (D, E, H, I, K, L, M, O, P, Q, S), Mann–Whitney test (F), ordinary one-way ANOVA testing (B), *P<0.05; **P<0.01; ***P<0.001; ****P<0.0001; ns, not significant. Scale bars: 10 μm.

### Somatic lysosomes are less degradative with neuronal aging

The number, size, and LAMP1 mean intensity of somatic LAMP1^+^ Els were analyzed in 21 and 28 DIV neurons using the ICY Bioimage analysis spot detector (de Chaumont et al., 2012). The number of somatic LAMP1^+^ Els remained constant between 21 and 28 DIV (Fig.1 C, D), whereas their size increased by 14%, and LAMP1^+^ mean intensity increased by 55% in aged somas (Fig.1 C, E, F), suggesting an up-regulation of endolysosomes with neuronal aging.

To determine if aging altered somatic LAMP1^+^ LEs, we co-stained neurons with the cation-independent mannose-6-phosphate receptor (CI-M6PR) that transports lysosomal hydrolases from the trans-Golgi network to endosomes. Importantly, CI-M6PR recycles from LEs, not being present in lysosomes (Ghosh et al., 2003). We found that nearly 20% of somatic LAMP1^+^ Els were positive for CI-M6PR in mature neurons, suggesting that one-fifth of somatic Els correspond to LEs. The percentage of LEs increased to nearly 30% in aged somas. In addition, CI-M6PR mean intensity *per* LAMP1^+^ LEs increased two-fold in aged somas (Fig.1 G, H, I), suggesting LEs accumulate with neuronal aging.

To determine if the somatic LAMP1^+^ lysosomes changed with neuronal aging, we co-stained neurons with Cathepsin D, the main lysosomal hydrolase, present both in degradative and non-degradative lysosomes (Bright et al., 2016). We found that nearly 15% of somatic LAMP1^+^ Els were positive for Cat D in mature neurons, which increased to 21% in aged neurons, suggesting that only a fraction of ELs are lysosomes or a suboptimal Cat D detection (Fig.1 J, K). Moreover, the size of LAMP1^+^ Cat D^+^ lysosomes increased with aging by 21% (Fig.1 J, L). We also found that Cat D accumulates in aged somatic lysosomes since the intensity of Cat D *per* lysosome increased by nearly 85% (Fig.1 J, M), suggesting a change in Cat D lifetime, like LAMP1.

To identify the degradative lysosomes, we used the lysosomal substrate DQ-BSA, cleaved by lysosomal proteases emitting bright fluorescence proportional to the lysosomal activity (Marwaha and Sharma, 2017). Notably, the DQ-BSA mean intensity *per* lysosome was brightly visible in the soma of 21 DIV neurons. Moreover, we found that 40% of somatic LAMP1^+^ Els accumulated DQ-BSA (Fig. 1 N, O), indicating that nearly half of LAMP1^+^ compartments represent degradative lysosomes, a higher percentage than the 15% detected with Cat D. In aged somas, the number of LAMP1^+^ DQ-BSA^+^ lysosomes was similar to mature somas. However, the DQ-BSA mean intensity was reduced by 50% (Fig. 1 N, O, P), indicating that the lysosomes are less degradative with neuronal aging. Moreover, the size of LAMP1^+^ DQ-BSA^+^ lysosomes was reduced by 12% in the aged soma, suggesting that the aged degradative lysosomes are smaller (Fig. 1 N, Q) in contrast with the enlarged LAMP1^+^ Els (Fig. 1 C, E).

We were intrigued by the reduced lysosomal degradative capacity in the aged somas, given the increased lysosomal hydrolase, Cat D. Therefore, we used the fluorescent acidotropic probe, LysoTracker (Barral et al., 2022), to determine whether the acidification of ELs was optimal for degradative activity. We observed a 60% reduction in LysoTracker labeling in the soma of 28 DIV neurons (Fig. 1 R, S), suggesting that the acidification of ELs is defective with aging, which may account for the reduced lysosomal degradation.

Overall, we establish that neuronal aging reduces the degradative capacity of somatic lysosomes, likely due to lysosome deacidification, leading to the accumulation and enlargement of LEs in the soma.

### Neuritic endolysosomes accumulate distally in aged neurons

Since LAMP1^+^ ELs concentrate in the proximal region of neurites in immature neurons (Yap et al., 2018), we analyzed whether neuritic LAMP1^+^ ELs positioning changed with the distance to the soma in mature and aged neurons. For that, we analyzed the number of LAMP1^+^ ELs in three neuritic regions starting from the soma: proximal region, 0–25 μm; medial region, 25–50 μm; and distal region, 50–75 μm. In mature neurons, 42 % of LAMP1^+^ ELs are in the neuritic proximal region, decreasing to 34 % and 24 % in the medial and distal regions of 21 DIV neurons, respectively (Fig. 2 A, B). Compared to mature neurites, the decline in aged neurites was less notorious, with the number of LAMP1^+^ ELs significantly increasing by 17 % in the medial and 35 % in the distal neurites (Fig. 2 A, B), suggesting that ELs accumulate distally in aged neurons. Moreover, we found that although the LAMP1^+^ ELs size declined after the proximal region in mature and aged neurons (Fig. 2 A, C), the distal LAMP1^+^ ELs size increased by 20% in aged neurons, suggesting an enlargement of distal ELs with aging. Of note, similarly to the soma, the LAMP1 mean intensity increased by 51%, 57%, and 40% in proximal, medial, and distal aged neurites, respectively (Fig. 2 A, D). The increase in LAMP1 mean intensity per ELs was not due to higher LAMP1 levels in aged neurons (Fig. S1 B, C) or aged brain (Fig.S1 E, F), suggesting that LAMP1 might accumulate in ELs while diminishing its presence in its other cellular locations such as the plasma membrane with aging (Janvier and Bonifacino, 2005; Saftig and Klumperman, 2009).

**Figure 2.**
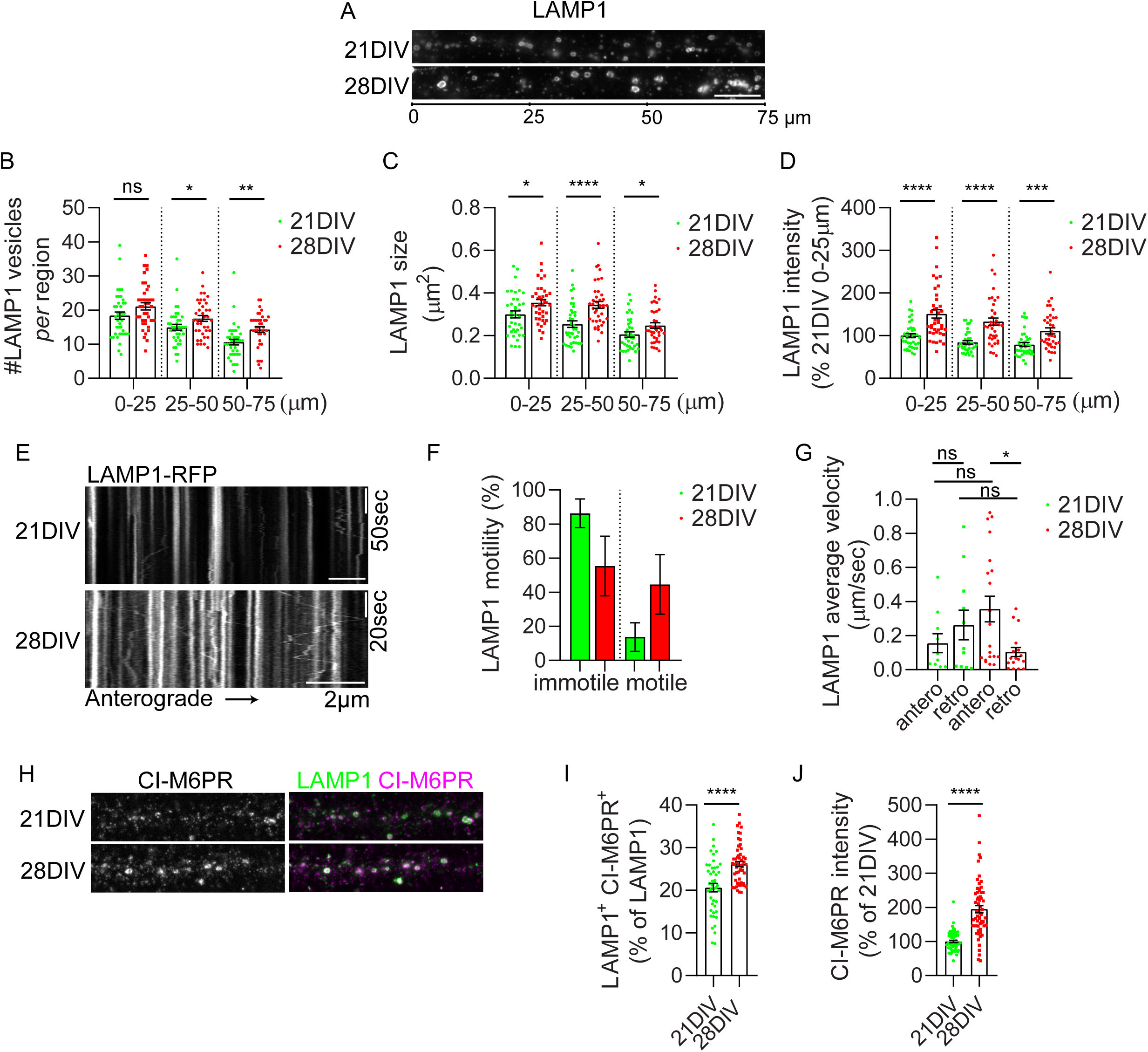
Neuritic endolysosomes accumulate distally in aged neurons. (A) LAMP1^+^ endolysosomes in neurites of 21 DIV and 28 DIV neurons. (B) Quantification of LAMP1^+^ endolysosome number *per* neurite in proximal (0-25µm), medial (25-50µm) and distal (50-75µm) region from the cell body (n=4, N neurites= 39). (C) Quantification of LAMP1^+^ endolysosome size in the proximal, medial, and distal regions of neurites (n=4, N neurites 0-25 µm = 38-40, N neurites 25-50 µm = 38-40, N neurites 50-75 µm = 37-38). (D) Quantification of LAMP1^+^ endolysosome mean intensity in the proximal, medial, and distal regions of neurites in the percentage of 21 DIV neurites proximal region (n=4, N neurites 0-25 µm = 39-40, N neurites 25-50 µm = 38-39, N neurites 50-75 µm = 37-39). (E) Kymographs of neurites from 21 and 28 DIV neurons transfected with LAMP1-RFP and GFP. (F) Quantification of LAMP1-RFP immotile and motile endolysosomes (n=2, N = 222-154 vesicles from 11-15 neurites). (G) Quantification of the average velocity of LAMP1-RFP endolysosomes (n=2, N = 26-62 motile vesicles from 11-15 neurites). (H) LAMP1^+^ (green) CI-M6PR^+^ (white; magenta) late-endosomes in neurites of 21 and 28 DIV neurons. (I) Quantification of the percentage of LAMP1^+^ endolysosomes colocalizing with CI-M6PR in neurites (n=3, N neurites=46-57). (J) Quantification of the mean intensity of CI-M6PR *per* LAMP1^+^ late-endosome in neurites in the percentage of 21 DIV neurons (n=3, N neurites = 61-62). Data are mean ± SEM. Statistical significance was determined by Mann–Whitney test (D, J) or unpaired t-test (B, C, I). One-Way ANOVA Holm-Sidak’s multiple comparison test (G). *P<0.05; **P<0.01; ***P<0.001; ****P<0.0001; ns, not significant. Scale bars: 10 μm

Because ELs accumulated in distal regions of aged neurites (Fig. 2 A, B), we hypothesized that aging could alter LAMP1^+^ ELs trafficking. We imaged LAMP1-RFP in live 21 and 28 DIV neurites to test our hypothesis for 2-3 min. We constructed kymographs and quantified the fraction of LAMP1-RFP^+^ ELs that were stationary, moved anterogradely, retrogradely, and their mean velocity (Pepperkok et al., 2005). We found that the fraction of motile LAMP1-RFP^+^ ELs tended to increase in 28 DIV neurons (Fig. 2 E, F). The analysis of the mean velocity of the motile LAMP1^+^ ELs revealed a higher mean velocity of LAMP1-RFP^+^ ELs in the anterograde direction in aged neurites, which could contribute to their distal accumulation since their retrograde mean velocity remained unaltered (Fig. 2 G). Interestingly, inhibition of synaptic increases ELs motility (Kulkarni et al., 2021), indicating that the increase in LAMP1^+^ ELs motility could be related to the aged neurons’ synapse loss (Burrinha et al., 2021).

Next, we investigated whether the neuritic ELs that accumulated in aged neurites were immature LEs or degradative lysosomes, hoping to clarify whether local degradation occurs distally near synapses or is restricted to the soma and proximal neurites (Farfel-Becker et al., 2019; Goo et al., 2017). We verified that the LEs fraction of ELs (20%) in neurites was similar to the soma in mature neurons. Interestingly, we detected a 30% increase in LAMP1^+^ LEs with a 90% increase in CI-M6PR content in aged neurites (Fig. 2 H, I, J). These results indicate that the LEs fraction of ELs increases significantly in aged neurites, like in the somas (Fig. 1 G, H, I).

Overall, enlarged ELs accumulated distally in aged neurites, linked to transport defects.

### Distal neuritic aged lysosomes accumulate Cat D but are less degradative

To determine if LAMP1^+^ ELs present in neurites were degradative lysosomes affected by neuronal aging, we co-stained neurons with Cat D and DQ-BSA and observed LAMP1^+^ Cat D^+^ DQ-BSA^+^ lysosomes along mature and aged neurites (Fig. 3 A, yellow arrows). In intensity line profiles of neurites, it is noticeable that the Cat D signal was brightest in the LAMP1^+^ lysosomes in aged neurites, especially in the proximal region (Fig. 3 B). In contrast, the DQ-BSA signal was reduced along aged neurites, often in the same lysosomes that accumulated Cat D (Fig. 3 C).

**Figure 3.**
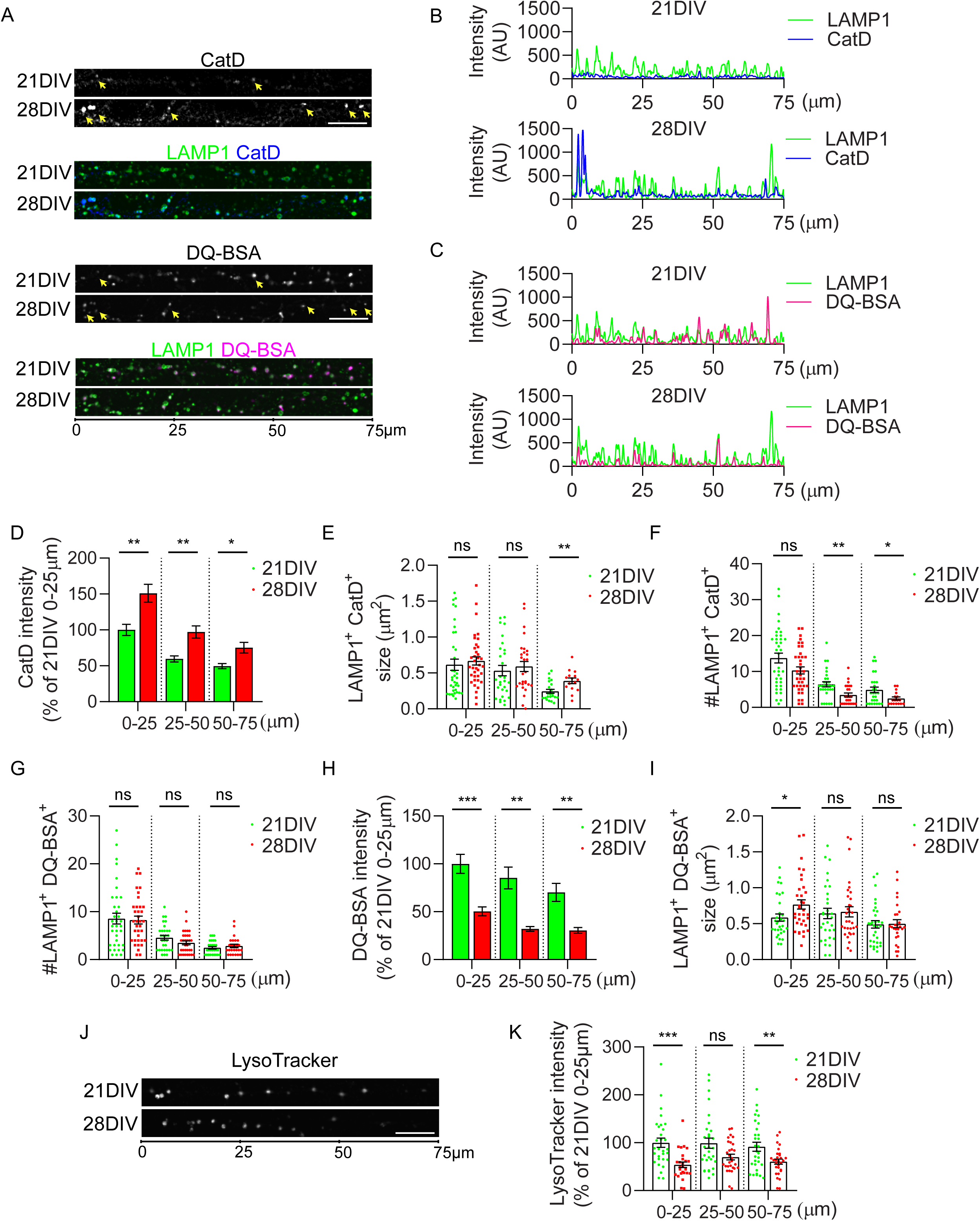
Distal lysosomes accumulate Cat D but are less degradative. (A) Cat D^+^ (white; blue), DQ-BSA^+^ (white; magenta), and LAMP1^+^ (green) lysosomes along 75µm of 21 and 28DIV neurites, displayed after background subtraction. (B) Line scan along the representative image in A to visualize the extent of peak coincidence of LAMP1 (green) and Cat D (blue) in 21 and 28 DIV neurons. (C) Line scan along the representative image in A to visualize the extent of peak coincidence of LAMP1 (green) and DQ-BSA (magenta) in 21 and 28 DIV neurons. (D) Quantification of the mean intensity of Cat D *per* LAMP1^+^ lysosomes in the proximal, medial, and distal regions of neurites in the percentage of 21 DIV neurites proximal region (n=3, N neurites 0-25 µm = 36-39, N neurites 25-50 µm = 38-39, N neurites 50-75 µm = 38-39). (E) Quantification of LAMP1^+^ Cat D^+^ lysosome size in the proximal, medial, and distal regions of 21 and 28 DIV neurites (n=3, N neurites 0-25 µm=36-37, N neurites 25-50 µm=28-29, N neurites 50-75 µm=15-22). (F) Quantification of LAMP1^+^ Cat D^+^ lysosome number in the proximal, medial, and distal regions of 21 DIV and 28 DIV neurites (n=3, N neurites 0-25 µm=37-38, N neurites 25-50 µm=25-30, N neurites 50-75 µm=17-27). (G) Quantification of LAMP1^+^ DQ-BSA^+^ lysosome number in the proximal, medial, and distal regions of 21 and 28 DIV neurites (n=3, N neurites 0-25 µm=36, N neurites 25-50 µm=30-32, N neurites 50-75 µm=25-28). (H) Quantification of the mean intensity of DQ-BSA *per* LAMP1^+^ lysosome in the proximal, medial, and distal regions of neurites in the percentage of 21 DIV neurites proximal region (n=3, N neurites 0-25 µm = 31-39, N neurites 25-50 µm = 31-39, N neurites 50-75 µm = 33-37). (I) Quantification of LAMP1^+^ DQ-BSA^+^ lysosome size in the proximal, medial, and distal regions of 21 DIV and 28 DIV neurites (n=3, N neurites 0-25 µm=32-35, N neurites 25-50 µm=31-32, N neurites 50-75 µm=27-31). (J) LysoTracker puncta in the proximal, medial, and distal regions of 21 and 28DIV neurites. (K) Quantification of the mean intensity of LysoTracker puncta in proximal, medial, and distal regions of neurites in the percentage of 21 DIV neurites proximal region (n=3, N neurites 0-25 µm= 27-30, N neurites 25-50 µm= 29-30, N neurites 50-75 µm= 29-30). Data are mean ± SEM. Statistical significance was determined by Mann–Whitney test (D, E, F, G, H, I, K). *P<0.05; **P<0.01; ***P<0.001; ns, not significant. Scale bars: 10 μm.

Quantification confirmed a striking increase in the Cat D intensity *per* LAMP1^+^ lysosomes in aged neurites (Fig. 3 D). The accumulation of Cat D translated into a significant increase in the size of LAMP1^+^ Cat D^+^ in distal aged neurites (Fig. 3 E), while their number also decreased in the medial regions of aged neurites (Fig. 3 F). This accumulation of Cat D in LAMP1^+^ lysosomes might relate to changes in trafficking since the levels of Cat D (mature and immature (Pro-Cat D)) were unaltered in aged neurons, despite a tendency for increased levels of mature Cat D (Fig. S1 B, D).

Moreover, quantification revealed that 50% of proximal LAMP1^+^ ELs accumulate DQ-BSA in mature neurites, reducing to 28% and 18% in medial and distal neurites, respectively. Notably, half of the LAMP1^+^ DQ-BSA^+^ lysosomes are in the medial/distal regions of mature neurites (Fig. 3 A, G), supporting that local degradation can occur more distally than in immature neurons (Yap et al., 2018). In aged neurites, the most striking difference was the reduction in DQ–BSA mean intensity in each lysosome along neurites, reaching a 60% reduction (Fig. 3 H). The size of LAMP1^+^ DQ-BSA^+^ lysosomes increased proximally in aged neurites (Fig. 3 A, I), while the number of LAMP1^+^ DQ-BSA^+^ lysosomes was unaltered (Fig. 3 G). Therefore, lysosome degradative capacity is significantly compromised in aged neurites, especially distally, which could explain the accumulation of LAMP1^+^ LEs. Of note, given that the degradative lysosomes (LAMP1^+^DQ-BSA^+^) are unaltered, a decrease in non-degradative lysosomes (LAMP1^+^Cat D^+^) may account for the reduction of LAMP1^+^Cat D^+^ lysosomes.

Since we did not detect a lack of lysosomal hydrolases that could explain the reduced degradative capacity of lysosomes in aged neurons, we evaluated their acidity using LysoTracker (Barral et al., 2022). LysoTracker brightly stained vesicles present along 21 DIV neurites, correlating with the accumulation of DQ-BSA in LAMP1^+^ lysosomes along neurites, consistent with acidic and degradative lysosomes in mature neurites, even distally (Fig. 3 J). In aged neurites, the intensity of LysoTracker was significantly reduced by 46% in the proximal and 34% in the distal regions (Fig. 3 J, K). The reduced LysoTracker and DQ-BSA mean intensity in aged neuritic lysosomes indicate a reduction in acidification underlying the reduced degradative activity of lysosomes in aged neurites, like in aged somas, despite the accumulation of Cat D.

### Enlarged endolysosomes accumulate at synapses in the aged brain

We evaluated LAMP1 changes in the prefrontal cortex (PFC) and hippocampus, specifically in the C*ornu Ammonis* 1 and 3 (CA1 and CA3) brain regions of adult wild-type mice (6 months old; 6M) and aged wild-type mice (18 months old; 18M). We found that, similarly to *in vitro* aged neurons, LAMP1^+^ ELs size was increased by 56% in CA1 and 27% in PFC but not in the CA3 region of the aged brain (Fig. 4 A, B). LAMP1 levels were unaltered, as shown by immunohistochemistry (IHC) quantification (Fig. 4 A, C) as well as by western blot (Fig. S1 E, F) in the aged brain, as previously reported (Huang et al., 2020).

**Figure 4.**
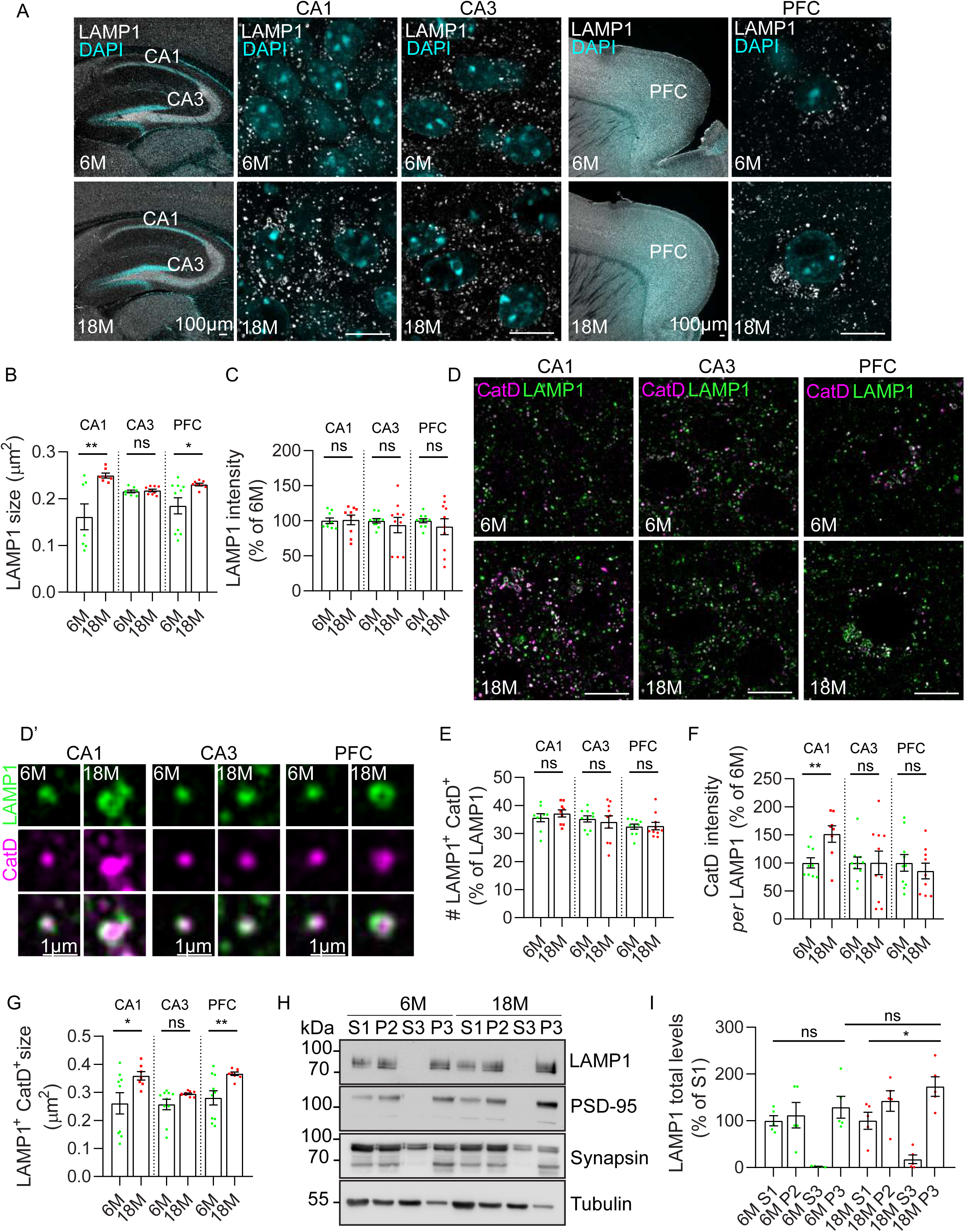
Endolysosomes are enlarged and accumulate at synapses in the aged brain. (A) LAMP1^+^ endolysosomes (white) in the hippocampus (CA1 and CA3) and prefrontal cortex (PFC) of 6-and-18-month-old mice brains, displayed after background subtraction. DAPI is in blue. (B) Quantification of LAMP1^+^ endolysosome size in CA1, CA3, and PFC of 6-and-18-month-old mice brains (n=3, N CA1 images = 7-8, CA3 images = 8-9, N PFC images = 8). (C) Quantification of the mean intensity of LAMP1 *per* endolysosome in CA1, CA3, and PFC of 6- and-18-month-old mice brains (n=3, N CA1 images = 9, CA3 images = 10, N PFC images = 10). (D) Cat D^+^ (magenta) LAMP1^+^ (green) lysosomes in CA1, CA3, and PFC of 6-and-18-month-old mice brains, displayed after background subtraction. Magnified lysosomes in CA1, CA3, and PFC of 6-and-18-month-old mice brains are shown in D’. (E) Quantification of the percentage of LAMP1^+^ Cat D^+^ lysosomes in CA1, CA3 and PFC regions (n=3, N CA1 images = 9, N CA3 images = 10, N PFC images = 10). (F) Quantification of the mean intensity of Cat D *per* LAMP1^+^ lysosome in CA1, CA3 and PFC of 6- and-18-month-old mice brains (n=3, N CA1 images = 8-9, CA3 images = 10, N PFC images = 9-10). (G) Quantification of LAMP1^+^ Cat D^+^ lysosome size in CA1, CA3 and PFC regions (n=3, N CA1 images = 7-9, N CA3 images = 9-10, N PFC images = 8-10). (H) Western blot analysis of LAMP1, PSD-95, synapsin, and tubulin as a loading control in post-nuclear supernatant (S1), crude synaptosomal fraction (P2), crude synaptic vesicle fraction (S3), and synaptosomal membrane fraction (P3) from 6-month-old and 18-month-old of brain homogenates levels. (I) Quantification of LAMP1 in the different brain fractions in the percentage of S1 (n=5). Data are mean ± SEM. Statistical significance was determined by unpaired t-test (B, C, E, F, G, I) *P<0.05; **P<0.01; ns, not significant. Scale bars: 10 μm.

To determine if lysosomes changed in the aged brain, we evaluated LAMP1^+^ Cat D^+^ lysosomes by IHC. We detected Cat D in nearly 30% of LAMP1^+^ ELs in 6M and 18M brains (Fig. 4 D, E). Like aged neurons, the intensity of Cat D *per* lysosome was 50% higher in the aged CA1 region (Fig. 4 D, D’, F), and mature Cat D levels tended to increase in the aged brain (Fig. S1 E, G). This Cat D increase in lysosomes is conserved in the aged rat and human brain and likely precedes the early AD accumulation of Cat D (Cataldo et al., 1995; Nakanishi et al., 1997). Notably, the Cat D lysosomes were 37% and 30% larger in aged CA1 and PFC regions, respectively (Fig. 4 D, D’, G), like aged somatic Cat D lysosomes (Fig. 1 L)

Since we found LAMP1^+^ ELs in a higher number in distal neurites of 28 DIV neurons, closer to synapses (Fig. 2 D, E), we evaluated LAMP1 levels in synaptosomes of the aged brain compared to the adult brain. Interestingly, LAMP1 was enriched by 73% in the synaptosomal fraction of the aged brain (Fig. 4 H, I). These results support that ELs accumulate distally close to synapses with aging *in vitro* and *in vivo*.

### Lysosome dysfunction contributes to the aging-dependent loss of synapses

Since lysosomes are essential for maintaining synapses (Farfel-Becker et al., 2019; Goo et al., 2017; Lund et al., 2021; Padamsey et al., 2017) and we found that lysosomes are dysfunctional in aged neurons, we decided to investigate if lysosomal dysfunction caused synapse loss in aged neurons.

We analyzed the ELs’ proximity to synapses by co-transfection of LAMP1-RFP and GFP to highlight the dendritic morphology and spines in our mature and aged neurons (Fig. 5 A). We detected LAMP1-RFP^+^ ELs in or close to dendritic spines, supporting similar observations in younger (16 DIV) neurons (Goo et al., 2017). To restore aged lysosomes, we used the mucolipin transient receptor potential channel 1 (TRPML1) agonist, mucolipin synthetic agonist 1 (ML-SA1, 20 μM, 16h), that by inducing lysosomal calcium efflux, reduces ELs pH in neurons (Bae et al., 2014; Hui et al., 2019) and increases protease activity (Xia et al., 2020). In non-neuronal cells, ML-SA1 treatment has been described to reduce the number and size of LAMP1^+^ ELs (Cao et al., 2017) but also to increase endo-lysosomal biogenesis by upregulating LAMP1 expression among other lysosomal genes (Zhang et al., 2016). Indeed, we found that treating aged neurons with ML-SA1 increased the acidification of aged lysosomes since the number of acidic vesicles positive for LysoTracker augmented proximally and distally, where the intensity of LysoTracker also increased (Fig. 5 B, C, D). The degradative capacity of aged lysosomes increased significantly with the ML-SA1 treatment, as evidenced by the augmented DQ-BSA mean intensity in the proximal region of neurites (Fig. 5 E, F) and by the larger number of similar-sized DQ-BSA^+^ lysosomes distally (Fig. 5 E, G, H). ML-SA1 treatment seems to improve lysosomes without broadly affecting LAMP1 ELs’ number and size (Fig. 5 E, I, J), suggesting that LAMP1^+^ LE must be largely unaffected by ML-SA1. Of note, ML-SA1 treatment increased LAMP1 mean intensity (Fig. 5 E, K), suggesting that, in neurons, ML-SA1 may lead to the upregulation of LAMP1 expression without affecting ELs biogenesis. Next, we assessed whether ML-SA1 treatment could reduce lipofuscin accumulation in lysosomes by improving lysosomes’ degradative capacity. ML-SA1 treatment decreased by 50% the number of lipofuscin^+^ lysosomes in the aged somas (Fig. 5 L, M), suggesting that improving lysosome function reduces the accumulation of this cellular aging hallmark.

**Figure 5.**
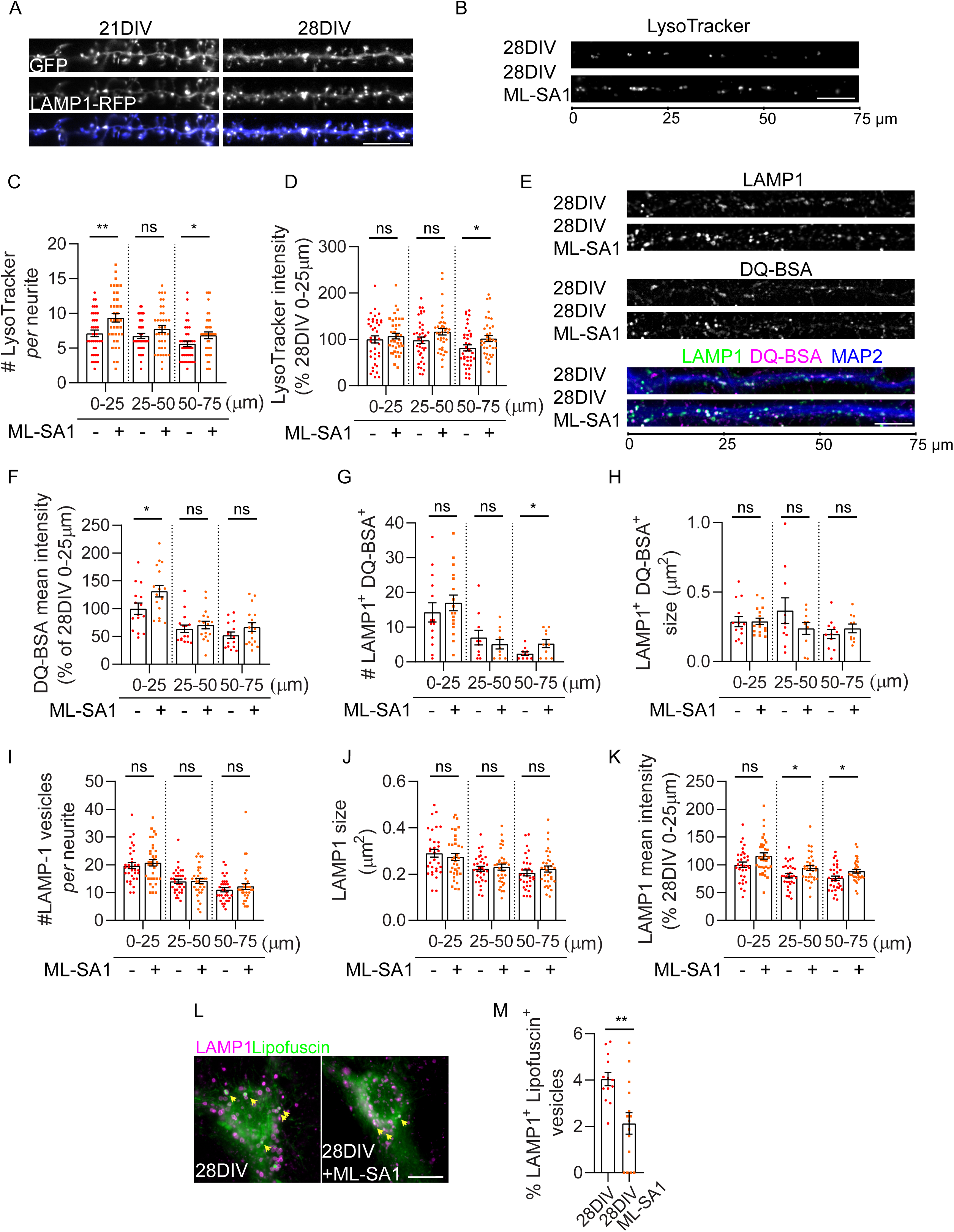
ML-SA1 increases lysosomal degradation and acidification of aged lysosomes. (A) Representative images of LAMP1-RFP in dendrites from 21 and 28 DIV neurons highlighted by GFP. (B) LysoTracker puncta in the proximal (0-25µm), medial (25-50µm), and distal (50-75µm) regions of neurites (28 DIV and 28 DIV ML-SA1). LysoTracker levels were adjusted at 28 DIV. (C) Quantification of LysoTracker puncta number in the proximal, medial, and distal regions of 28 DIV and 28 DIV neurites treated with ML-SA1 (n=4, N neurites 0-25 µm=38-39, N neurites 25-50 µm=38-39, N neurites 50-75 µm=38-39). (D) Quantification of LysoTracker puncta mean intensity in proximal, medial, and distal regions of neurites, in the percentage of 28 DIV neurites proximal region (n=4, N neurites 0-25 µm= 38-39, N neurites 25-50 µm= 38-39, N neurites 50-75 µm= 37-39). (E) DQ-BSA^+^ (magenta; white) LAMP1^+^ (green; white) lysosomes along 75um neurites (28 DIV and 28 DIV ML-SA1), immunolabeled with MAP2 (blue), displayed after background subtraction. (F) Quantification of the mean intensity of DQ-BSA *per* LAMP1^+^ lysosome in the proximal, medial, and distal regions of neurites in the percentage of 28 DIV neurites proximal region (n=2, N neurites 0-25 µm= 15-17, N neurites 25-50 µm= 15-17, N neurites 50-75 µm=15-17). (G) Quantification of LAMP1^+^ DQ-BSA^+^ lysosome number in the proximal, medial, and distal regions of 28 DIV and 28 DIV neurites treated with ML-SA1 (n=2, N neurites 0-25 µm=15-17, N neurites 25-50 µm=10, N neurites 50-75 µm=10). (H) Quantification of LAMP1^+^ DQ-BSA^+^ lysosome size in the proximal, medial, and distal regions of 28 DIV and 28 DIV neurites treated with ML-SA1 (n=2, N neurites 0-25 µm=14-17, N neurites 25-50 µm=10, N neurites 50-75 µm=10-11). (I) Quantification of LAMP1^+^ endolysosome number in the proximal, medial, and distal region of neurites (n=4, N neurites= 32-34). (J) Quantification of LAMP1^+^ endolysosome size in the proximal, medial, and distal regions of neurites (n=4, N neurites 0-25 µm= 32-34, N neurites 25-50 µm= 32-34, N neurites 50-75 µm=31-34). (K) Quantification of the mean intensity of LAMP1^+^ endolysosomes in the proximal, medial, and distal regions of neurites, in the percentage of 28 DIV neurites proximal region (n=4, N neurites 0-25 µm= 32-34, N neurites 25-50 µm= 32-33, N neurites 50-75 µm=32). (L) Lipofuscin (green) and LAMP1 (magenta) localization in the cell body of neurons (28 DIV and 28 DIV ML-SA1). (M) Quantification of the percentage of LAMP1^+^ lysosomes containing lipofuscin (n=3, N cell body=13-15). Data are mean ± SEM. Statistical significance was determined by Mann–Whitney test (C, D, G, H, I, K) or unpaired t-test (F, J, M). *P<0.05; **P<0.01; ns, not significant. Scale bars: 10 μm.

To determine if restoring lysosome degradative capacity was relevant for neuronal function, we assessed the impact of ML-SA1 treatment on the synapse loss that we previously reported in aged neurons (Burrinha et al., 2021). As previously, we quantified synapse density by measuring the number of juxtapositions between a presynaptic marker (vGluT1) and the postsynaptic marker (PSD-95) along dendrites (Fig. 6 A, B; (Burrinha et al., 2021)). Similarly to our previous observations, synapse density was reduced by 40 % in aged neurons (Fig. 6 A, B). Remarkably, ML-SA1 treatment of aged neurons increased synapse density by 34 % (Fig. 6 A, B). We also analyzed the pre- and postsynaptic compartments separately. We found that the ML-SA1 treatment rescued the decrease in PSD-95 puncta size and mean intensity without altering vGluT1 puncta size (Fig. 6 A, C-F), indicating that lysosome dysfunction with aging has a more relevant impact on postsynaptic function.

**Figure 6.**
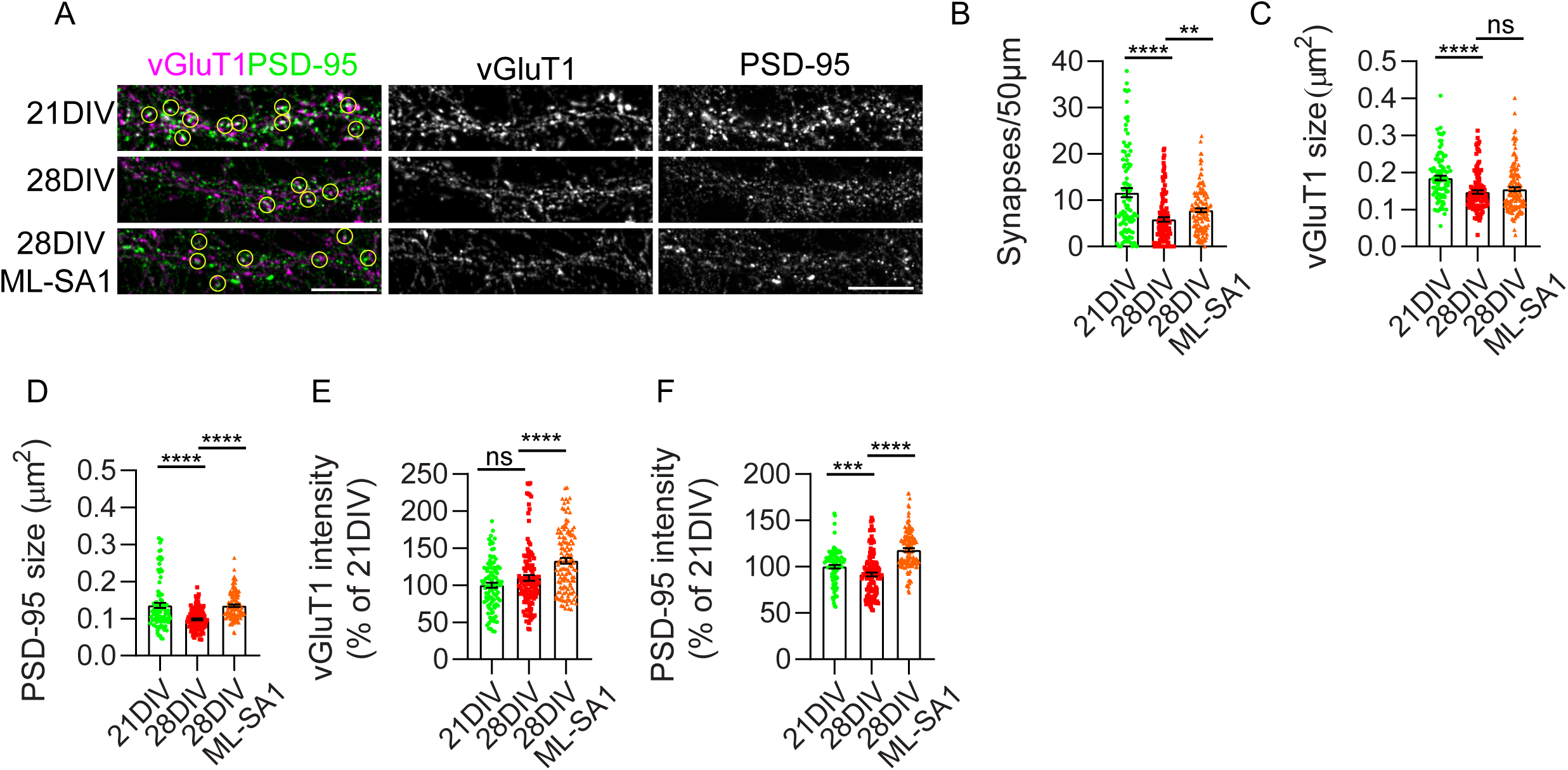
Lysosome dysfunction contributes to the aging-dependent loss of synapses. (A) PSD-95 (green; white) and vGluT1 (magenta; white) in 21 DIV, 28 DIV, and 28 DIV neurites treated with ML-SA1, displayed after background subtraction. Yellow rings indicate synapses. (B) Quantification of the number of synapses *per* 50 μm of neurites in neurons with 21 DIV, 28 DIV, and 28 DIV treated with ML-SA1 (n=3; N 21 DIV=96 neurites; N 28 DIV=123 neurites; N 28 DIV ML-SA1=126 neurites). (C) Quantification of vGluT1 size *per* neurite in neurons with 21 DIV, 28 DIV, and 28 DIV treated with ML-SA1 (n=3; N 21 DIV=96 neurites; N 28 DIV=125 neurites; N 28 DIV ML-SA1=128 neurites). (D) Quantification of PSD-95 size *per* neurite in neurons with 21 DIV, 28 DIV, and 28 DIV treated with ML-SA1 (n=3; N 21 DIV=96 neurites; N 28 DIV=127 neurites; N 28 DIV ML-SA1=128 neurites). (E) Quantification of vGluT1 mean intensity in neurons with 21 DIV, 28 DIV, and 28 DIV treated with ML-SA1 (n =3, N 21 DIV=96 neurites; N2 8 DIV= 123 neurites; N 28 DIV ML-SA1= 128 neurites). (F) Quantification of PSD-95 mean intensity in neurons with 21 DIV, 28 DIV, and 28 DIV treated with ML-SA1 (n =3, N 21 DIV= 96 neurites; N 28 DIV=127 neurites; N 28 DIV ML-SA1=128 neurites). Data are mean ± SEM. Statistical significance was determined by Mann–Whitney test (B, C, D, E, F). **P<0.01; ***P<0.001; ****P<0.0001; ns, not significant. Scale bars: 10 μm.

Overall, our data suggest that restoring aged lysosomes’ acidification improves lysosomal degradative capacity in aged neurons, rescuing age-dependent synapse loss and delaying cellular aging.

### Inducing lysosome dysfunction recapitulates age-dependent loss of synapses

To determine if the induction of lysosome dysfunction would recapitulate the age-dependent synapse loss, we treated 21 DIV neurons with the lysosomotropic agent chloroquine (CQN). CQN alkalinizes the lysosomal lumen (Fedele and Proud, 2020), causing endo-lysosomal enlargement and reduced degradation (Fedele and Proud, 2020; Ponsford et al., 2021).

We evaluated the impact of CQN treatment (20 μM, 16h) on lysosome number, size, positioning, degradation capacity (LAMP1^+^DQ-BSA^+^), and LAMP1^+^ ELs’ number, size, and positioning along neurites. CQN reduced the number of lysosomes accumulating DQ-BSA in proximal neurites by 40% (Fig. 7 A, B). Moreover, in the degradative lysosomes, the DQ-BSA accumulation was reduced by 33% (Fig. 7 C). This decreased degradative activity likely underlies the observed enlargement of DQ-BSA^+^ lysosomes in proximal and medial regions of neurites (Fig. 7 D). Together, these data suggest that lysosomal degradation decreases with CQN treatment. In addition, CQN treatment recapitulated aged neurons’ endo-lysosomal enlargement, with the size of LAMP1^+^ ELs increasing by more than 65% along neurites (Fig. 7 E). LAMP1^+^ ELs decreased by 20% in the medial region (Fig. 7 F), and LAMP1 intensity tended to increase along neurites (Fig. 7 G). These results indicate that the CQN treatment is sufficient to cause lysosome enlargement and degradative dysfunction proximally but not distally, partially recapitulating the neuritic aging phenotype.

**Figure 7.**
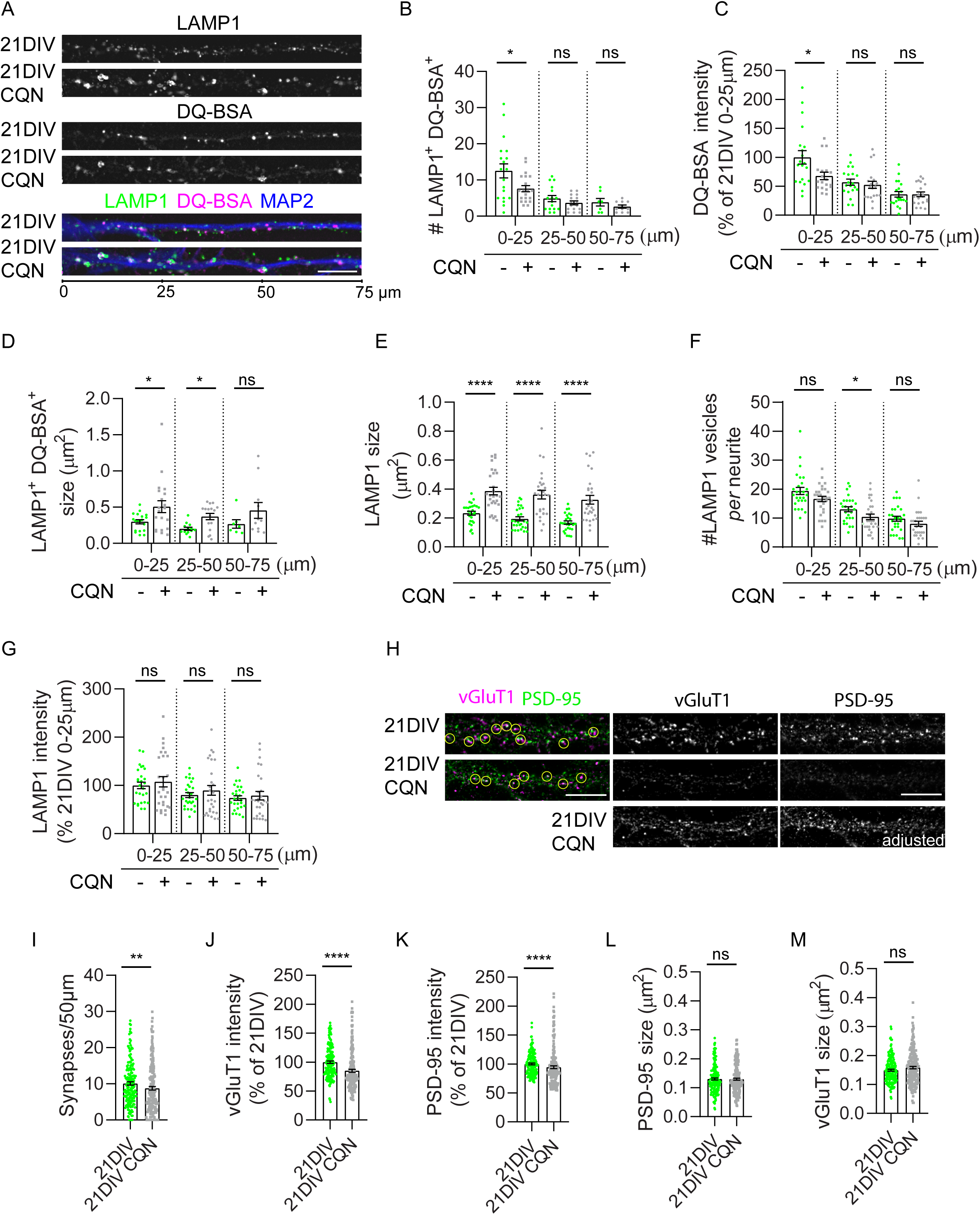
Inducing lysosome dysfunction recapitulates age-dependent loss of synapses. (A) DQ-BSA^+^ (white; magenta) LAMP1^+^(white; green) lysosomes along 75µm neurites (21 DIV and 21 DIV CQN), immunolabeled with MAP2 (blue), displayed after background subtraction. (B) Quantification of LAMP1^+^ DQ-BSA^+^ lysosome number in the proximal (0-25µm), medial (25-50µm), and distal (50-75µm) regions of 21 DIV and 21 DIV neurites treated with CQN (n=3, N neurites 0-25 µm=19-20, N neurites 25-50 µm=16, N neurites 50-75 µm=7-11). (C) Quantification of the mean intensity of DQ-BSA *per* LAMP1^+^ lysosome in the proximal, medial, and distal regions of neurites in the percentage of 21 DIV neurites proximal region (n=3, N neurites 0-25 µm=19-20, N neurites 25-50 µm=20, N lysosomes 50-75 µm=20). (D) Quantification of LAMP1^+^ DQ-BSA^+^ lysosome size in the proximal, medial, and distal regions of 21 DIV and 21 DIV neurites treated with CQN (n=3, N neurites 0-25 µm=19-20, N neurites 25-50 µm=13-16, N neurites 50-75 µm=7-11). (E) Quantification of LAMP1^+^ endolysosome size in the proximal, medial, and distal regions of neurites (n=4, N neurites 0-25 µm=27-28, N neurites 25-50 µm=27-28, N neurites 50-75 µm=27-28). (F) Quantification of LAMP1^+^ endolysosome number in proximal, medial, and distal regions of neurites (n=4, N neurites= 27-28). (G) Quantification of LAMP1 mean intensity *per* endolysosome in the proximal, medial, and distal regions of neurites in the percentage of 21 DIV neurites proximal region (n=4, N neurites 0-25 µm=27-28, N neurites 25-50 µm=27-28, N neurites 50-75 µm=27-28). (H) PSD-95 (green; white) and vGluT1 (magenta; white) in 21 DIV and 21 DIV neurites treated with CQN, displayed after background subtraction. Yellow rings indicate synapses. (I) Quantification of the number of synapses *per* 50 μm of neurites in neurons with 21 DIV and 21 DIV treated with CQN (n=3; N 21 DIV=167 neurites; N 21 DIV CQN=198 neurites). (J) Quantification of the mean intensity of vGluT1 in neurites with 21 DIV and 21 DIV treated with CQN (n=3; N 21 DIV=173 neurites; N 21 DIV CQN=200 neurites). (K) Quantification of the mean intensity of PSD-95 in neurites with 21 DIV and 21 DIV treated with CQN (n=3; N 21 DIV=173 neurites; N 21 DIV CQN=197 neurites). (L) Quantification of PSD-95 size in neurites with 21 DIV and 21 DIV treated with CQN (n=3; N 21 DIV=173 neurites; N 21 DIV CQN=197 neurites). (M) Quantification of vGluT1 size in neurites with 21 DIV and 21 DIV treated with CQN (n=3; N 21 DIV=173 neurites; N 21 DIV CQN=200 neurites). Data are mean ± SEM. Statistical significance was determined by Mann–Whitney test (C, D, E, F, I, J, K, L, M) or unpaired t-test (B, G). *P<0.05; **P<0.01; ****P<0.0001; ns, not significant. Scale bars: 10 μm.

Finally, we determined that CQN treatment reduced synapse density in mature neurons (Fig. 7 H, I), with a reduced mean intensity of vGluT1 and PSD-95 (Fig. 7 H, J, K) while not affecting the size of vGluT1 and PSD-95 (Fig. 5 H, L, M). Our data agree with previous reports of synapse disruption induced by other lysosomotropic agents (Kanju et al., 2007). Overall, our results implicate lysosomal dysfunction as a causal mechanism of age-dependent synapse loss.

## Discussion

Our recent discovery that intracellular Aβ accumulation only partially accounts for neuronal aging-dependent synapse decline (Burrinha et al., 2021) prompted us to investigate the other mechanism(s) involved. Since lysosome dysfunction is a hallmark of cellular aging (López-Otín et al., 2013), lysosome function is relevant for synapses (Farfel-Becker et al., 2019; Goo et al., 2017; Padamsey et al., 2017), and we showed that lysosomes accumulate lipofuscin with neuronal aging (Burrinha et al., 2021), we set out to investigate lysosome dysfunction with neuronal aging and whether it contributes to synapse loss. We characterized endolysosomes (ELs) (late endosomes and lysosomes) in mature and aged neurons. In mature neurons, we found degradative lysosomes in neurites, even distally. In aged neurons, we discovered that enlarged endolysosomes accumulate distally, likely due to increased anterograde transport. Significantly, lysosome degradative capacity decreases throughout aged neurons. This lysosomal impairment correlates with reduced ELs acidification but not with the increased content in Cat D. We demonstrate that restoring lysosome acidification and function improves synapses in aged neurons. In contrast, deacidifying lysosomes in mature neurons recapitulated aged lysosome dysfunction and synapse loss. Therefore, we propose that neuronal aging-dependent-lysosome dysfunction can trigger synapse loss.

### Characterization of the lysosome in mature neurons

We found that the LAMP1^+^ endo-lysosomal system in mature neurons comprises 20% of late-endosomes (LAMP1^+^CI-M6PR^+^) and 40% of lysosomes (LAMP1^+^DQ-BSA^+^). The 40% unidentified LAMP1^+^ compartments may correspond to post-Golgi secretory carriers, like in axons (Lie et al., 2021). Lysosomes have been localized in the neuronal soma and proximal neurites (Cai et al., 2010; Cheng et al., 2018; Ferguson, 2019; Gowrishankar et al., 2015; Lee et al., 2011; Tammineni et al., 2017; Yap et al., 2018), but degradation can occur near synapses (Goo et al., 2017; Jin et al., 2018). We found a larger endo-lysosomal area in neurites than in the soma of mature neurons (21 DIV). Strikingly, about one-fourth of LAMP1^+^ ELs in mature distal neurites are degradative lysosomes (Fig. 2 E and 3 G), more than in immature neurons (9 DIV) (Yap et al., 2018). This distal increase of degradative lysosomes in neurites may relate to our neurons being synaptically more mature, influencing lysosome trafficking and positioning (Goo et al., 2017; van Bommel et al., 2019). Of note, even in the soma, only half of the LAMP1^+^ ELs are lysosomes. Thus, LAMP1 alone does not identify lysosomes (Barral et al., 2022). Our results support that local lysosomal degradation occurs distally in mature neurites.

### Neuronal aging causes lysosomal dysfunction

Our findings in neurons aged *in vitro* and *in vivo* support that neuronal aging alters the endo-lysosomal system (Fig. 1, 2, 3, and 4). We found late-endosomes (LAMP1^+^CI-M6PR^+^) and non-degradative lysosomes (LAMP1^+^Cat D^+^) increased and enlarged, while the degradative lysosomes (LAMP1^+^DQ-BSA^+^) were less active in aged neurons. Moreover, we established that deacidification of aged lysosomes likely underlies their diminished degradative capacity and ELs enlargement, similarly to other aged cells (Mai et al., 2019; Truschel et al., 2018). The enlargement of lysosomes observed in neurons aged *in vitro* was confirmed in the brain of aged mice, with the aged CA1 region being the most affected (Fig. 4). Interestingly, loss of functional synapses occurs mainly in the aged CA1 region (Barnes et al., 1992; Barnes et al., 2000).

The acidification of the lysosome, a pre-requisite to activate most lysosomal hydrolases, is achieved by the ATP-dependent proton pump, the vacuolar ATPase (v-ATPase) but also depends on the efflux of cations by channels such as TRPML1, which mediates lysosomal calcium release (Colacurcio and Nixon, 2016). Although lysosome deacidification had not been shown in aged neurons, it was described in aged *S. cerevisiae* (Ruckenstuhl et al., 2014). Additionally, reducing the activity of v-ATPase has been shown to cause cognitive impairment in *Drosophila Melanogaster* and *Mus Musculus* (Dubos et al., 2015). It would be essential to determine whether v-ATPase and TRPML1 decrease with neuronal aging.

Despite unaltered total levels, the increase in LAMP1, Cat D, and CI-M6PR mean intensity *per* ELs was another striking phenotype. Surprisingly, despite LAMP1 being the most used ELs marker, its increased mean intensity in ELs had not been previously described in neurons of the aged brain. Regarding Cat D, mean intensity, total levels, and lifetime increase in the aged rat brain (Kluever et al., 2022; Nakanishi et al., 1994; Nakanishi et al., 1997). Interestingly, LAMP1 and Cat D increased in blood exosomes ten years prior to the development of mild cognitive impairment (Goetzl et al., 2015).

The increased LAMP1, CI-M6PR, and Cat D content of ELs may relate to augmented trafficking to endolysosomes. In agreement, endocytosis is upregulated with neuronal aging (Alsaqati et al., 2018; Blanpied et al., 2003; Burrinha et al., 2021). The increase in CI-M6PR may also contribute to Cat D transport to endosomes, explaining Cat D lysosomal enrichment (Fox et al., 1992; Ghosh et al., 2003; Guay et al., 2000; Zaidi et al., 2008). Moreover, Cat D maturation seems unaffected by lysosomal dysfunction with aging (Fig. S1).

Since the transcription factor EB (TFEB), which activates LAMP1 and Cat D expression, is reduced in the aged mouse brain (Wang et al., 2021), it is unlikely to underlie the increase in LAMP1 and Cat D in lysosomes. Little is known about LAMP1 and Cat D degradation and whether their degradation is affected by aging. It may not be since restoring lysosomal degradation in aged neurons did not reduce the levels of LAMP1 (Fig. 5), suggesting that other mechanisms regulate LAMP1 lysosomal levels.

In addition, we found that the number of ELs increased mainly in aged distal neurites *in vitro* and aged synaptosomes *in vivo*. This phenotype might be explained by the reduced lysosome acidification distally in aged neurites (Fig. 3) since deacidification increases lysosome movement towards the cell periphery in fibroblasts (Heuser, 1989) and into neurites (Parton et al., 1991). However, neither the increased acidification of ELs in aged neurons (ML-SA1 treatment) corrected the endo-lysosomal positioning, nor the deacidification of ELs in mature neurons (CQN treatment) recapitulated the distal accumulation of ELs (Fig. 5, 7).

Alternatively, the positioning of ELs may depend instead on transport (Bonifacino and Neefjes, 2017; Heuser, 1989; Johnson et al., 2016; Parton et al., 1991), either on increased anterograde transport to distal neurites for local degradation (Farfel-Becker et al., 2019; Farfel-Becker et al., 2020) and/or on decreased retrograde transport to the soma (Kimura et al., 2012). Interestingly, aged ELs moved faster anterogradely than retrogradely, explaining the distal ELs accumulation. Alternatively, we envision two underlying mechanisms 1) reduced diffusion due to the enlargement of aged ELs since lysosome size influences diffusion but not their active transport (Bandyopadhyay et al., 2014; de Araujo et al., 2020), 2) less tethering of ELs to the actin cytoskeleton due to the reduction in spines of aged neurites (Goo et al., 2017; van Bommel et al., 2019) or to changes in the actin cytoskeleton in aged neurons (Alexander Mack et al., 2016).

### TRPML1 activation restores synapse decline in aged neurons

We found that the functional changes observed in aged ELs contribute to synapse decline since treatment with an agonist of TRPML1 (ML-SA1) increased acidification (Bae et al., 2014), the degradative capacity of lysosomes, and increased synapses in aged neurons (Fig. 5, 6). Remarkably, neuronal aging reduced PSD-95 size, as reported in mature spines of aging learning-impaired rat brains (Nicholson et al., 2004). Curiously, ML-SA1 treatment significantly increased the PSD-95 size, which suggests that lysosome exocytosis might be increased (Di Paola et al., 2018; Medina et al., 2011), releasing cathepsins that may contribute to spine remodeling via degradation of the extracellular matrix (Padamsey et al., 2017). Another possibility is that ML-SA1 might restore the local lysosomal degradation of synaptic membrane proteins such as AMPA receptors (Ehlers, 2000; Goo et al., 2017). The number and size of ELs were not changed with ML-SA1 treatment of aged neurons, ruling out a contribution of lysosome biogenesis to the synapse rescue (Sardiello et al., 2009).

Importantly, the ML-SA1 treatment decreased lipofuscin accumulation in aged neurons (Fig. 5) (Burrinha et al., 2021), suggesting that lipofuscin is a consequence and not a cause of lysosome dysfunction with neuronal aging.

### Induction of lysosome dysfunction partially mimics the age-dependent synapse loss

We found that CQN treatment induced lysosome dysfunction, with reduced degradative activity, deacidification, and enlarged ELs, in agreement with previous studies (Fedele and Proud, 2020; Ponsford et al., 2021). Interestingly, longer *in vivo* induction of lysosome dysfunction leads to an increase in the number of lysosomes and lipofuscin accumulation (Ivy et al. 1989; Gray and Woulfe 2005; Sohal and Wolfe 1986; Sohal and Brunk 1989; Bednarski et al. 1997; Ivy et al. 1984). Importantly, we discovered that CQN treatment compromises synapses in mature neurons (Fig. 7). Our results indicate that lysosome deacidification through CQN treatment is sufficient to cause lysosome and synapse dysfunction recapitulating neuronal aging. It remains unclear how lysosome function maintains synapses, whether via degradation of synaptic cargo, autophagosomes containing damaged organelles or protein aggregates, or other lysosomal functions such as metabolic sensing and membrane repair.

### The impact of dysfunctional lysosomes on aging, longevity, and neurodegenerative diseases

Our data support a causal role for lysosome dysfunction in age-dependent synapse decline in the brain. Future research is needed to dissect whether the lysosome buffering of cellular calcium (Ballabio and Bonifacino, 2020) is altered with aging since calcium increases in the aged brain (Nikoletopoulou and Tavernarakis, 2012), and it may impact neuronal excitability and synaptic plasticity. Moreover, it will be essential to determine the brain’s regional vulnerabilities since not all hippocampal and cortical regions suffer a decline with aging at the same rate (Raz and Rodrigue, 2006).

The time-dependent slow disruption in ELs function and trafficking with neuronal aging might potentiate the initiation of age-related neurodegenerative disorders such as AD. Indeed, impaired lysosomal activity is associated with dysfunctional clearance of autophagic substrates (Boland et al., 2018; Bordi et al., 2016) and neuritic swellings in LOAD brains (Lee et al., 2011). The lysosome dysfunction with aging broadens the lysosomes as a hotspot in AD, adding to the lysosome dysfunction induced by mutations in presenilins causative of familial AD (Cataldo et al., 2004; Coen et al., 2012)

## Conclusion

Lysosomal function defects may arise during neuronal aging, reducing its degradative ability and contributing to synapse aging. The gradual loss of function of lysosomes with neuronal aging may not completely arrest the endo-lysosomal system. Instead, lysosome dysfunction may generate vulnerability to risk factors that could trigger age-related neurodegenerative diseases such as AD. Therefore, understanding the endo-lysosomal system in neuronal aging will highlight the early mechanisms of age-related synapse loss and potentiate the development of therapeutic strategies to delay the onset of neurodegenerative disorders.

## Limitations of the study

We used primary embryonic cortical neurons from wild-type mice and aged them for 28 days *in vitro* to investigate mechanisms of neuronal aging. This cellular system of neuronal aging *in vitro* allows for imaging at high resolution, gene manipulation, and dissection of cellular and molecular mechanisms of synapse decline in a relatively short time, which is not possible with *in vivo* models of aging. Moreover, the morphology and physiology of mouse neurons are similar to human neurons. However, this model considers the cell-intrinsic defects in neurons with aging but has the limitation of not representing the influence of cell-extrinsic alterations on the systemic environment of the aging organism (Pluvinage and Wyss-Coray, 2020). Still, brain aging hallmarks, such as lipofuscin accumulation and synapse loss (Gray and Woulfe, 2005; Morrison and Baxter, 2012; Samson and Barnes, 2013), occur in our aged *in vitro* cultures. Therefore, with the *in vivo* validation in the physiologically aged brain, despite measuring lysosome acidification and activity *in vivo* not being technically possible, we consider the present model a suitable tool to study how synapse impairment develops in an age-dependent manner.

## Author contributions

Conceptualization and project management: CGA. Funding acquisition and supervision: CGA; Investigation: TB, CGA, CC; Methodology and formal analysis: TB, CGA, CC; visualization and writing: TB, CGA.

## Acknowledgments

We thank Dr. Stephanie Miserey-Lenkei (Institute Curie), Dr. Duarte Barral (NOVA Medical School), Dr. Otília Vieira (NOVA Medical School) for the gift of antibodies and reagents. We thank Dr. André Marques, Dr. Cristina Escrevente, Dr. Catarina Perdigão, and Liliana Lopes for helpful discussions. We thank L. Almeida for excel macros. We thank the lab members for helpful discussions and for the critical reading of the manuscript. We thank Dr. Duarte Barral (NOVA Medical School) and Dr. Otília (NOVA Medical School) for the critical reading of the manuscript. We thank Dr. Ana Farinho (NOVA Medical School Histology Facility), Dr. S. Marques (NOVA Medical School Animal Facility) and Dr. T. Pereira (NOVA Medical School Microscopy Platform). This project has received funding from Marie Curie Integration Grant (PCIG-GA-2012-334366-trafficinAD; Marie Curie Actions, EC); iNOVA4Health—UID/Multi/04462/2019, a program financially supported by Fundação para a Ciência e Tecnologia (FCT)/ Ministério da Educação e Ciência, through national funds and co-funded by FEDER under the PT2020 Partnership Agreement); Maratona da Saúde 2016; Investigator FCT (IF/00998/2012, FCT); CEECIND/00410/2017 financed by FCT; ALZ AARG-19-618007 Alzheimer’s Association; from the research infrastructure PPBI-POCI-01-0145-FEDER-022122, co-financed by FCT (Portugal) and Lisboa2020, under the PORTUGAL2020 agreement (European Regional Development Fund); the European Union’s Horizon 2020 research and innovation program under grant agreement No 811087 (Lysocil). TB has been the recipient of an FCT doctoral fellowship (SFRH/BD/131513/2017).

## Declaration of interests

The authors declare no competing interests.

## Materials and Methods

### Animals

All animal procedures were performed according to EU recommendations and were approved by the Ethical Committee, the NMS – Universidade Nova de Lisboa ethical committee (142/2021/CEFCM), and the national DGAV (Direção-Geral da Alimentação e Veterinária; 0421/000/000/2013).

### Cell culture

Primary neuronal cultures from *Mus musculus* were prepared as previously (Almeida et al., 2005) from cortices of embryonic day 16 (E16) wild-type females and male BALB/c mice. Briefly, E16 brain tissue was dissociated by trypsinization and trituration in Dulbecco’s medium Eagle medium (DMEM) with 10% fetal bovine serum (Heat-Inactivated FBS, Thermo Fisher Scientific). Dissociated neurons were plated in DMEM with 10% FBS on poly-D-lysine (Sigma-Aldrich)-coated six-well plates (1×10^5^ cells/cm2) and glass coverslips (2.6×10^4^ cells/cm2). After 3-16 h, the medium was substituted for Neurobasal medium supplemented with B27, GlutaMAX, and penicillin/streptomycin (all from Life Technologies) at 37°C in 5% CO2. Neurons were maintained for 28 days *in vitro* (DIV) without changing or adding a new medium. We only used healthy cultures that presented minimal glia. When indicated, TRPML1 was activated by 16 h treatment with 20µM of ML-SA1 (Abcam, ab144622), lysosome acidification and enzymatic function were inhibited by 16 h treatment with 20 μM of Chloroquine (Sigma, C6628), and DMSO (0.1%; solvent; PanReac AppliChem) was used as a control.

### cDNA expression

For cDNA expression (LAMP1-RFP and GFP), neurons (12 DIV) were transiently transfected with 4μg of cDNA (live-cell imaging) or 1μg of cDNA (immunofluorescence) with Lipofectamine 2000 (Life Technologies) for 5 min and were analyzed after 21 DIV or 28 DIV.

### Antibodies

The following primary antibodies were used: anti-MAP2 mAb [Sigma-Aldrich, M4403, 1:500 (IF, IHC)]; anti-tubulin mAb [Tu-20, Millipore, MAB1637, 1:10,000 (WB)]; anti-LAMP1 mAb [CD107a, BD Pharmingen, 553792; 1:200 (IF); 1:500 (IHC); 1:1000 (WB)]; anti-PSD-95 mAb [Merck Millipore, 04-1066, 1:200 (IF)]; anti-vGluT1 mAb [Merck Millipore, MAB5502, 1:200 (IF)]; anti-CathepsinD mAb [ab75852, Abcam, EPR3057Y, 1:1000 (WB)]; anti-CathepsinD mAb [Milipore, 06-467, 1:200 (IF), 1:500 (IHC)]; anti-CI-M6PR mAb [ab2733, Abcam, 2G11; 1:200 (IF)]; anti-Synapsin I pAb [Abcam, ab8, 1:1000 (WB)]. The secondary antibodies used were conjugated to Alexa Fluor 488, 555 and 647 (Molecular Probes), or to horseradish peroxidase (HRP, Bio-Rad).

### Immunofluorescence labeling

Immunofluorescence was performed as previously (Almeida et al., 2005; Ubelmann et al., 2017). Briefly, cultured primary neurons were fixed at 21 and 28 DIV with 4% paraformaldehyde/4% sucrose in PBS for 20 min, permeabilized with 0.1% saponin in PBS for 1 h, and blocked in 2% FBS/1% bovine serum albumin (BSA)/0.1% saponin in PBS for 1 h at room temperature before antibody incubation using a standard procedure. For PSD-95 and vGluT1 immunolabeling, permeabilization was performed with 0.3% Triton X-100 in PBS for 5 min at room temperature. DAPI (Sigma, D9542) was added to the secondary antibody solution. Coverslips were mounted using Fluoromount-G (Southern Biotechnology, Birmingham, AL, USA).

### Image acquisition

Epifluorescence microscopy was carried out on an upright microscope Z2 (Zeiss) equipped with a 63× NA-1.4 oil immersion objective and an AxioCam MRm charged-coupled device (CCD) camera (Zeiss). Live imaging of LysoTracker was performed on a Revolution xD (Andor) confocal spinning-disc system coupled to an Eclipse Ti-E microscope (Nikon), whereas LAMP1—RFP and GFP was performed with AiryScan - LSM 980 confocal microscope (AiryScan 2, Zeiss). During imaging, the temperature was maintained at 37°C using a temperature-controlled incubator. Confocal microscopy was performed with the AiryScan-LSM 980 confocal microscope (AiryScan 2, Zeiss). Samples were imaged in two dimensions, with the focus plane chosen based on signal sharpness and best signal-to-noise ratio, in parallel, and using identical acquisition parameters for direct comparison.

### Immunoblotting

Cell lysates were prepared using modified RIPA buffer [50 mM Tris-HCl (pH 7.4), 1% NP-40, 0.25% sodium deoxycholate, 150 mM NaCl, 1 mM EGTA, and 0.1% SDS, with protease inhibitor cocktail (PIC)] as described (Burrinha et al., 2021). Proteins separated by 12% Tris-glycine SDS–PAGE were transferred to nitrocellulose membranes (GE Healthcare) and processed for immunoblotting using an ECL Prime kit (GE Healthcare). Images of immunoblots were captured using a ChemiDoc Gel Imaging System (Bio-Rad) within the linear range and quantified by densitometry using the ‘Analyse gels’ function in ImageJ.

### Lysosome assays

To measure lysosomal degradative activity, neurons were incubated overnight at 37°C in a complete medium supplemented with 10 µg/ml DQ™ Red BSA (Invitrogen, D-12051) essentially as described (Sharifi et al., 2010). Briefly, prior to fixation, neurons were washed with PBS to remove excessive DQ-BSA. Subsequently, neurons were fixed, and the immunofluorescence labeling was performed as described previously. To assess endo-lysosomal acidification, neurons were incubated for 10min at 37°C in a complete medium supplemented, or in a pre-warmed live imaging medium (120 mM NaCl, 3 mM KCl, 2 mM CaCl2, 2 mM MgCl2, 10 mM glucose, 10 mM HEPES), with 100nM LysoTracker™ Red DND-99 (Invitrogen, L7528). Neurons were imaged live or washed with PBS, fixed with 4% paraformaldehyde/4% sucrose in PBS for 20 min, and imaged immediately after mounting.

### Quantitative analyses

Image analyses were carried out using ImageJ (imagej.nih.gov/ij), Fiji (Fiji.sc) or ICY (icy.bioimageanalysis.org; (de Chaumont et al., 2012). For the quantification of LAMP1^+^ area per subcellular compartment (Cell body vs Neurites), in each individual neuron ICY ‘region of interest (ROI) - 2D ROI – Polygon or Area was used to outline 1 ROI - cell body and 3 ROIs -Neurites (Fig. 1). LAMP1^+^ endolysosomes were segmented using the ICY “Spot Detector” plugin. The sum of the LAMP1^+^ endolysosomes area was represented *per* ROI - cell body and *per* ROIs - neurites *per* individual neuron.

For the quantification of CI-M6PR enrichment per LAMP1^+^ endolysosome (Fig. 1, 2), the ICY’ spot detector’ plugin was used to create a ROI - endolysosome, the average of mean fluorescence of each channel *per* ROI - endolysosome was obtained with ICY ROI *per* Cell body or neurite.

For the quantification of the late-endosome number, the number of colocalizations of LAMP1 with CI-M6PR was obtained using the ICY “Colocalization Studio - SODA” plugin (statistical object distance analysis; SODA) (Lagache et al., 2018), considering the distance between objects less than 5 pixels.

For the quantification of the number, size, and mean fluorescence intensity of LAMP1^+^ endolysosomes, LAMP1^+^ Cat D^+^ lysosomes, and LAMP1^+^ DQ-BSA^+^ lysosomes in the cell body (Fig. 1), as well as the % of LAMP1+ endolysosomes containing lipofuscin in aged vs aged treated with ML-SA1 neurons (Fig. 5), the cell body was outlined using the ‘polygon selection’ tool on ImageJ, and the “ComDet v.0.5.5” plugin used for segmentation, colocalization, and the other measurements considering the max distance between objects less than 5 pixels.

For the quantification of LysoTracker mean fluorescence intensity (Fig. 1) *per* cell body, in live-cell imaging movies was quantified with the ‘measure’ function in the ROI - cell body outlined using Image J ‘Polygon selection’. The mean fluorescence intensity of LysoTracker was calculated as a percentage of 21 DIV upon background fluorescence subtraction.

For the quantification of LAMP1^+^ endolysosome number, size, and mean fluorescence intensity in neurites, the neurites that could be traced without interference from other processes were segmented using ImageJ “freehand lines” and “straighten”. Afterward, the ICY’ spot detector’ plugin was used on the selected neurite segments to detect LAMP1+ endolysosomes in three different neuritic sections, proximal (0–25 μm), medial (25–50 μm), and distal (50–75 μm) measured from the edge of the cell body (Fig. 2). For the quantification of Cat D or DQ-BSA enrichment *per* LAMP1^+^ lysosomes, upon LAMP1^+^ vesicle segmentation and the creation of ROIs using the ICY ‘spot detector’ plugin, the mean fluorescence *per* LAMP1 in each ROI was obtained using ICY “ROI statistics” *per* neurite. The mean fluorescence was calculated as a percentage of the mean intensity at the 21 DIV proximal region and upon background fluorescence subtraction (Fig. 3, 5, 7). To address the number and size of LAMP1^+^ Cat D^+^ and LAMP1^+^ DQ-BSA^+^ lysosomes, we filtered the vesicles with Cat D/DQ-BSA fluorescence intensity higher than the mean *per* neurite region (Fig. 3, 5, 7). To address the number and size of LAMP1^+^ Cat D^+^ lysosomes *in vivo*, we filtered the vesicles with Cat D fluorescence intensity higher than the mean *per* image (Fig. 4). The quantification of LysoTracker mean fluorescence intensity in neurites was done similarly to LAMP1^+^ endolysosomes in neurites (Fig. 3, 5).

For the quantification of LAMP1-RFP motility, kymographs were generated by the ImageJ plugin “Multiple Kymograph”, where traces of stationary or moving particles were marked, and the mean velocity of the moving particles was obtained using the macro “read velocities from tsp.” as previously described (Pepperkok et al., 2005)(Fig. 2).

For the quantification of LAMP1^+^ endolysosome size, mean fluorescence intensity, and Cat D enrichment *per* LAMP1 *in vivo* (Fig. 4), the ICY “Spot Detector” plugin as above was used to segment LAMP1 in a ROI largely corresponding to the neuronal somas based on the DAPI channel using the ICY “ROICreateFromChannel” tool. The mean fluorescence intensity of LAMP1 and Cat D *per* LAMP1 was divided by the mean intensity *per* image for each channel and calculated as a percentage of the 6-month-old mice, upon background fluorescence subtraction. To calculate the % of LAMP1^+^ Cat D^+^ lysosomes, we filtered the ones whose mean intensity was higher than the mean and calculated it as a percentage of the number of LAMP1^+^ lysosomes.

For the quantification of PSD-95 colocalization with vGluT1 (synapse) density *per* length of neurite (Fig. 6, 7), the number of colocalizing objects was obtained using the ICY ‘colocalizer’ with the “Spot detector” module protocol, as described (Burrinha et al., 2021). The density was calculated per length (Feret’s diameter) as indicated (Fig. 6, 7).

### Brain homogenates preparation

Frozen forebrains (including cortex and hippocampus) from 6-month-old (adult) and 18-month-old (aged) C57BL/6 mice were solubilized using a modified RIPA buffer. After 15 min on ice, brain lysates were further homogenized by sonication and centrifuged at 4°C for 30 min. Equivalent amounts of protein (40 μg), as determined by the BCA protein assay kit (Thermo Fisher Scientific), were mixed with sample buffer, heated at 95°C for 5 min, and separated by a 12% Tris-glycine SDS–PAGE.

### Brain synaptosomes preparation

Synaptosomes were prepared from forebrains (including cortex and hippocampus) from 6-month-old (adult) and 18-month-old (aged) C57BL/6 mice. All steps were performed at 4°C. Samples were homogenized with a pestle in ice-cold buffer 1 (0.32 M sucrose (Sigma); 10mM HEPES; 2 mL PIC (1X); 1mM EDTA, 1× complete protease inhibitor cocktail tablet; pH 7.4; diluted in ddH_2_O) using a 10 ml buffer per gram of tissue. The homogenate was centrifuged (10 min, 1000xg, 4°C) and the postnuclear supernatant (S1) was saved and centrifuged (15min, 10 000xg, 4°C) to generate a pellet that contains crude synaptosomes (P2). P2 was resuspended in 500 µl of buffer 1 and centrifuged (15min, 10 000xg, 4°C) generating the washed synaptosome fraction (P2’). P2’ was lysed by hypoosmotic shock in ice-cold ddH_2_O + 10mM HEPES, homogenized with a pestle, and let under head-over-heels rotation at 4°C for 30min to ensure complete lysis. P2’ was further centrifuged (30min, 21 000xg, 4°C) to generate a supernatant that contains crude synaptic vesicles (S3) and a pellet that contains synaptosomal membranes (P3). P3 was resuspended in 100 µl modified RIPA buffer [50 mM Tris-HCl (pH 7.4), 1% NP-40, 0.25% sodium deoxycholate, 150 mM NaCl, 1 mM EGTA, and 0.1% SDS, with protease inhibitor cocktail (PIC)].

### Brain immunohistochemistry

Histology was performed on fixed gelatin-embedded forebrains of the left-brain hemispheres of 6-month-old (adult) and 18-month-old (aged) BALB/c mice. Sagittal 40 μm sections were sliced on a microtome (Cryostat Leica CM3050 S). Sections were washed in 0.1 M phosphate buffer, permeabilized with 0.3% Triton X-100 in 0.1 M phosphate buffer for 10 min at room temperature, and blocked in 0.3% Triton X-100/5% BSA in 0.1 M phosphate buffer for 1 h at room temperature before primary antibody incubation in blocking solution overnight at 4°C. The secondary antibodies used were conjugated to Alexa Fluor 488, 555, and 647 (Molecular Probes). Coverslips were mounted using Fluoromount-G (Southern Biotechnology, Birmingham, AL, USA).

### Statistics

GraphPad Prism 8 was used for graphic representation of individual replicates with mean ± S.E.M. In most experiments, statistical analysis was performed in at least three independent experiments (n) as indicated in figure legends. The sample size (N) was determined based on pilot studies. Data were tested with the D’Agostino-Pearson omnibus normality test. For parametric and paired data, the paired t-test was applied; for parametric and unpaired data, the unpaired t-test was applied; for non-parametric and unpaired data, the Mann–Whitney test was applied; for nonparametric and multiple comparisons statistical analysis of data, one-way ANOVA on ranks with post-hoc Dunn’s testing was applied; for parametric and multiple comparisons statistical analysis of data, One-Way ANOVA Holm-Sidak’s multiple comparison test was applied. Outliers were removed with ROUT (Q = 1%) using GraphPad.

## Supplemental information title and legend

**Figure S1 – Aged lysosomes accumulate lipofuscin and LAMP1 and Cathepsin D levels do not change with aging *in vitro* and *in vivo*.**

(A) Lipofuscin (green) and LAMP1 (magenta) localization in the cell body of 21 DIV and 28 DIV neurons.

(B) Western blot analysis of LAMP1, pro-Cat D, mature-Cat D, and tubulin as a loading control in 21 DIV and 28 DIV neurons.

(C) Quantification of LAMP1 normalized to tubulin in 21 DIV and 28 DIV neurons (n=4).

(D) Quantification of pro-Cat D and mature-Cat D normalized to tubulin in 21 DIV and 28 DIV neurons (n=4).

(E) Western blot analysis of LAMP1, pro-Cat D, mature-Cat D, and tubulin as a loading control in mice brains (6-month-old - 6M, and 18-month-old – 18M).

(F) Quantification of LAMP1 normalized to tubulin in 6M and 18M mice brains (n=3-4).

(G) Quantification of pro-Cat D and mature-Cat D normalized to tubulin in 6M and 18M mice brains (n=3-4).

Data are mean ± SEM. Statistical significance was determined by Mann–Whitney test (C, D, F, G). ns, not significant. Scale bars: 10 μm.

## Notes

### Competing Interest Statement

The authors have declared no competing interest.

