## Supplementary figures and images for "Deacidification of distal lysosomes by neuronal aging drives synapse loss"

### Fig.S1

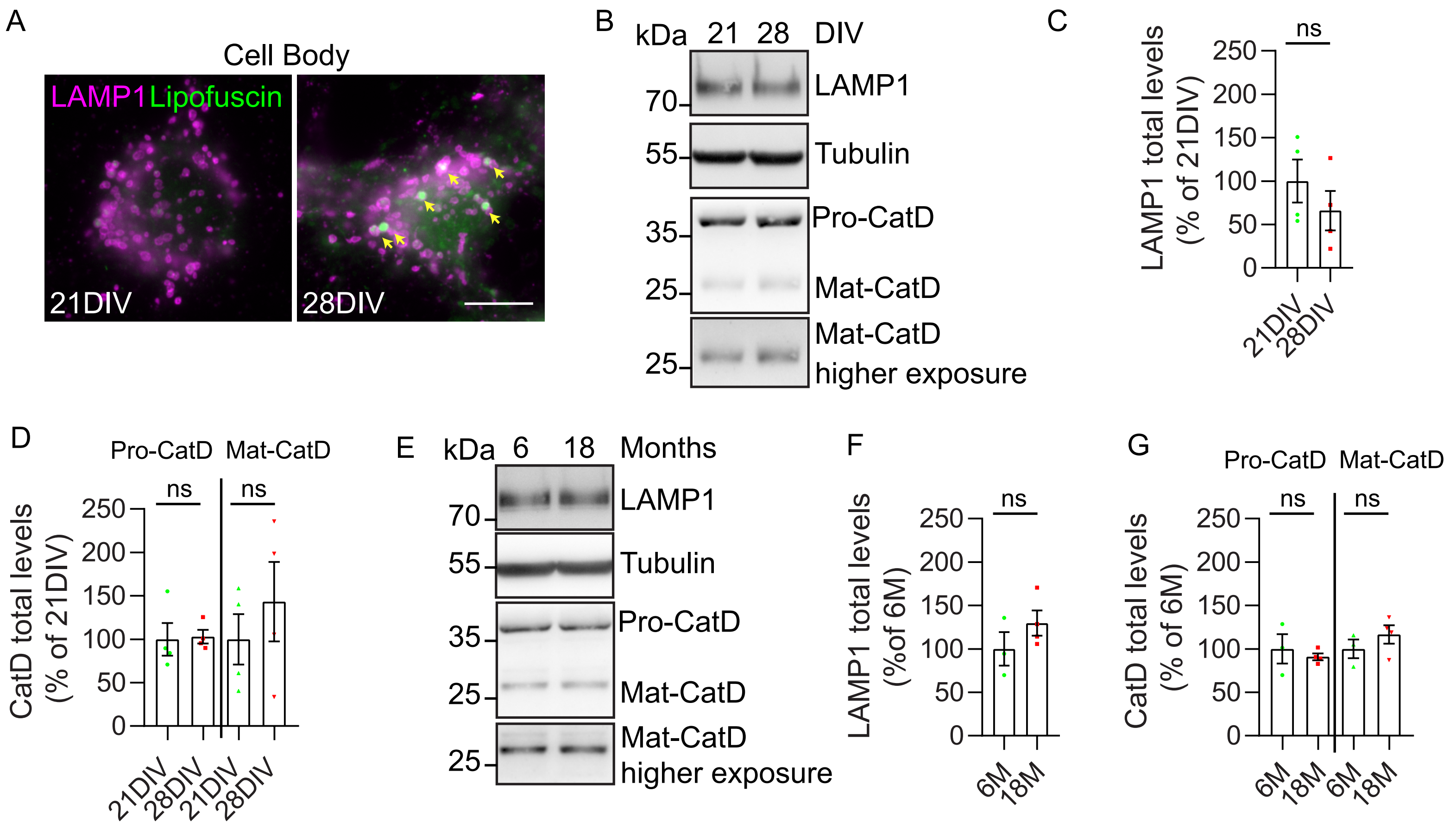
